# Itaconate promotes the differentiation of murine stress erythroid progenitors by increasing Nrf2 activity

**DOI:** 10.1101/2023.03.11.532211

**Authors:** Baiye Ruan, Yuanting Chen, Sara Trimidal, Imhoi Koo, Jingwei Cai, John Mcguigan, Molly A. Hall, Andrew D. Patterson, K. Sandeep Prabhu, Robert F. Paulson

## Abstract

Steady state erythropoiesis produces new erythrocytes at a constant rate to replace senescent erythrocytes removed in the spleen and liver. Inflammation caused by infection or tissue damage skews bone marrow hematopoiesis, increasing myelopoiesis at the expense of steady state erythropoiesis. To compensate for the loss of production, stress erythropoiesis is induced. Stress erythropoiesis is highly conserved between mouse and human but utilizes a different strategy than steady state erythropoiesis. Inflammatory signals promote the proliferation of immature stress erythroid progenitors (SEPS), which in response to erythropoietin and other signals transition to stress erythroid progenitors committed to differentiation. Here we show that TNFα dependent signaling increases ROS in SEPs during the proliferation stage, however, blocking ROS production impairs their later differentiation. In addition to TNFα, nitric oxide dependent signaling drives the proliferation of stress erythroid progenitors and production of nitric oxide must be decreased so that the progenitor cells can differentiate. As progenitor cells transition to differentiation, increased production of the anti-inflammatory metabolite itaconate activates Nfe2l2 or Nrf2, which inhibits Nos2 expression, leading to decreased nitric oxide production. Mutation of Irg1, the enzyme that catalyzes the production of itaconate, causes a delayed recovery from inflammatory anemia induced by heat killed *Brucella abortus*. Loss of itaconate-dependent activation of Nrf2 is rescued in vivo by IL-10, which leads to activation of Nrf2 and differentiation. These data show that the differentiation of stress erythroid progenitors relies on a switch to an anti-inflammatory metabolism and increased expression of pro-resolving cytokines.

**Key points:**

1. The transition to differentiation of stress erythroid progenitors requires anti-inflammatory signals.
2. The anti-inflammatory metabolite itaconate and IL-10 increase Nrf2 activity to promote the differentiation of stress erythroid progenitors.

## Introduction

Newly produced erythrocytes have a lifespan of approximately 120 days in humans and 45 days in mice, after which they are phagocytosed by resident macrophages in spleen and liver(^1^). To offset the turnover, new erythrocytes are constantly replenished by steady-state erythropoiesis. However, inflammation caused by infection or tissue damage disrupts this equilibrium. Pathogen associated molecular patterns or alarmins released from damaged tissues initiate inflammation via the production of inflammatory signaling molecules, such as nitric oxide (NO) and pro-inflammatory cytokines. Inflammatory signals shape the kinetics and routes of hematopoietic differentiation leading to increased production of myeloid cells at the expense of steady state erythropoiesis(^2–7^). Additionally, inflammation increases erythrophagocytosis which shortens the life span of erythrocytes(^8–11^). These demand-adapted changes in the blood system underpin a protective immune response to inflammatory insults, but they come with a cost as steady-state erythropoiesis is compromised. Given that effective oxygen transport requires adequate levels of erythrocytes, a compensatory stress erythropoiesis response is activated that utilizes inflammatory signals to produce erythrocytes and maintain homeostasis during inflammation(^12^).

Stress erythropoiesis is a stem cell-based tissue regeneration process that relies on the interaction between two major cellular components, stress erythroid progenitors (SEPs) and the monocyte-derived macrophage niche(^13^). Acting as an initiating signal, inflammation-induced erythrophagocytosis increases the production of chemokine CCL2 to recruit monocytes into the spleen where they interact with immature SEPs to form erythroblastic islands (EBIs)(^14^). In addition, erythrophagocytosis enhances the expression of inflammatory signals, such as TNF-α and NO, as well as the production of growth factors BMP4 and GDF15 in the niche(^15^). These secreted factors mediate extrinsic regulation on SEPs triggering the expansion of a large population of transient amplifying progenitors, while at same time they promote the development of niche monocytes into inflammatory macrophages(^15–18^). Increasing expression of erythropoietin (Epo) in the kidney, and the rise of serum Epo levels acts on macrophages leading to a phenotype switch from inflammatory to resolving macrophages(^19–21^). Aided by the switch of signals in the niche, immature SEPs commit to differentiation generating a bolus of new erythrocytes to reestablish homeostasis. Prostaglandin J2 (PGJ2) is a key lipid mediator of this transition response. Epo-induced production of PGJ2 activates PPARγ signaling, which decreases the expression of pro-proliferation factors (Wnt2b and Wnt8a) and increases the production of PGE2 which promotes the transition to SEP differentiation(^19^). This change in signals is like that observed when macrophages promote the resolution of inflammation by increasing the expression of IL-4, IL-10 and the anti-inflammatory metabolite itaconate(^22, 23^).

The tight spatiotemporal coordination between niche cells and SEPs suggest that they may utilize the same immunomodulatory molecules to regulate their co-development. Whereas the role of these molecules in macrophages is extensively studied, it remains undetermined how immunoregulatory metabolites and cytokines act on SEPs to affect their development. In this study, we applied integrated analysis of transcriptional and metabolic profiling on SEPs at different developmental stages. We show that transition of SEPs from expansion to differentiation is dependent on a switch of progenitor cell signaling from one dominated by inflammatory signals to one dominated by resolving signals. In the expansion stage, the increased TNF-α-dependent ROS production and iNOS-dependent NO production work in concert to drive the proliferation of immature SEPs. In contrast, the transition to SEP differentiation is marked by decreased inflammatory signals and increased resolving molecules like itaconate and IL-10. The anti-inflammatory effects of itaconate and IL-10 are mediated by the nuclear factor erythroid 2-related factor 2 (Nfe2l2 or Nrf2). Activation of Nrf2 resolves inflammatory signals in both SEPs and the niche, which reduces NO levels and alleviates the NO-mediated erythroid inhibition. These data provide a mechanistic basis of how SEP cell-fate transition is governed by immunomodulatory molecules to ensure effective erythroid regeneration.

## Methods

### Mice

Wild-type C57BL/6J, B6.SJL-Ptprca Pepcb/BoyJ (CD45.1) JAX stock #002014, B6.129X1-Nfe2l2tm1Ywk/J (Nrf2-/-) JAX stock #017009(^24^), B6.129P2-*Il10^tm1Cgn^*/J (IL-10-/-) JAX stock# 002251(^25^) and C57BL/6N-Acod1em1(IMPC)J/J (Irg1-/-) JAX stock #029340 mice were purchased from Jackson Laboratories. Mice of both sexes at age 8-16 weeks were used throughout this study. All experiments were approved by the Institutional Animal Care and Use Committee (IACUC) at the Pennsylvania State University.

### Statistics

GraphPad Prism and R were used for statistical analysis. Statistical significance between two groups was determined by two-tailed unpaired t test, except for the human culture experiment where paired t test was performed. Data with more than two groups were assessed for significance using one-way or two-way ANOVA followed by specific post hoc test as noted in the figure legends. Tukey’s test was used to make every possible pairwise comparison, whereas Dunnett’s correction was used to compare every group to a single control. Hematocrit levels were measured at different time points from same cohorts of mice, and data were analyzed by two-way repeated measures ANOVA followed by unpaired t test. Data are presented as mean ± SEM. Less than 0.05 of p value is considered as significant difference. n.s., p > 0.05; *, p < 0.05; **, p < 0.01; ***, p < 0.001.

### Data sharing statement

RNA-sequencing data have been deposited at NCBI GEO with accession number GSE190030. Metabolomics data have been deposited at NMDR (DOI: http://dx.doi.org/10.21228/M89402).

Additional methods are available in supplemental methods.

## Results

### Immature SEPs utilize inflammatory signal-dependent production of ROS for SEP expansion and maturation.

Growing bone marrow cells in stress erythropoiesis expansion media (SEEM) recapitulates the expansion of SEPs observed *in vivo*, whereas switching cultures to differentiation media (SEDM) mimics the transition of SEPs to differentiation(^15, 16, 18, 19, 26, 27^). We performed RNA-seq analysis to compare the transcriptome profile of SEPs isolated from SEEM and SEDM cultures. The expression of genes in proliferating SEPs were significantly enriched in pathways associated with inflammatory response, as well as NF-κB and pro-inflammatory cytokine-related signaling (Figure 1A-B). In contrast, these inflammatory signals were resolved in differentiating progenitors. We previously showed that the injection of zymosan increases the expression of pro-inflammatory cytokine TNF-α in the spleen, which functions to promote stress erythropoiesis(^15^). Consistent with this finding, we observed that proliferating SEPs had elevated TNF-α signaling (Figure 1A-B). The transient amplifying population of SEPs co-expresses Kit and Sca1, and they can be further divided into three subpopulations based on marker CD34 and CD133 (Figure S1A)(^18^). The addition of recombinant TNF-α protein into SEEM cultures increased the expansion of Kit^+^Sca1^+^ SEPs and facilitated the development of SEPs from Kit^+^Sca1^+^CD34^+^CD133^+^ to Kit^+^Sca1^+^CD34^-^CD133^-^ progenitor populations (Figure 1C). The latter population is more mature and more rapidly proliferative(^16^). These data build on our previous finding to show that immature SEPs utilize the inflammatory signal TNF-α for expansion and maturation. In immune cells, TNF-α has been known to promote the generation of mitochondrial reactive oxygen species (ROS) which regulates inflammation, immune cell function and proliferation(^28^). We found that proliferating SEPs also increased the production of mitochondrial ROS in response to TNF-α treatment (Figure 1D). Corresponding to elevated TNF-α signaling, ROS levels increased during SEP expansion (Figure 1E). However, this increase was compromised when SEEM cultures were supplemented with N-acetyl-l-cysteine (NAC), a ROS scavenger (Figure 1F). In contrast to the effect of TNF-α, NAC treatment blocks the expansion of Kit^+^Sca1^+^ SEPs (Figure 1C and 1F). These results demonstrate that TNF-α promotes SEP expansion by increasing ROS production.

**Figure 1.**
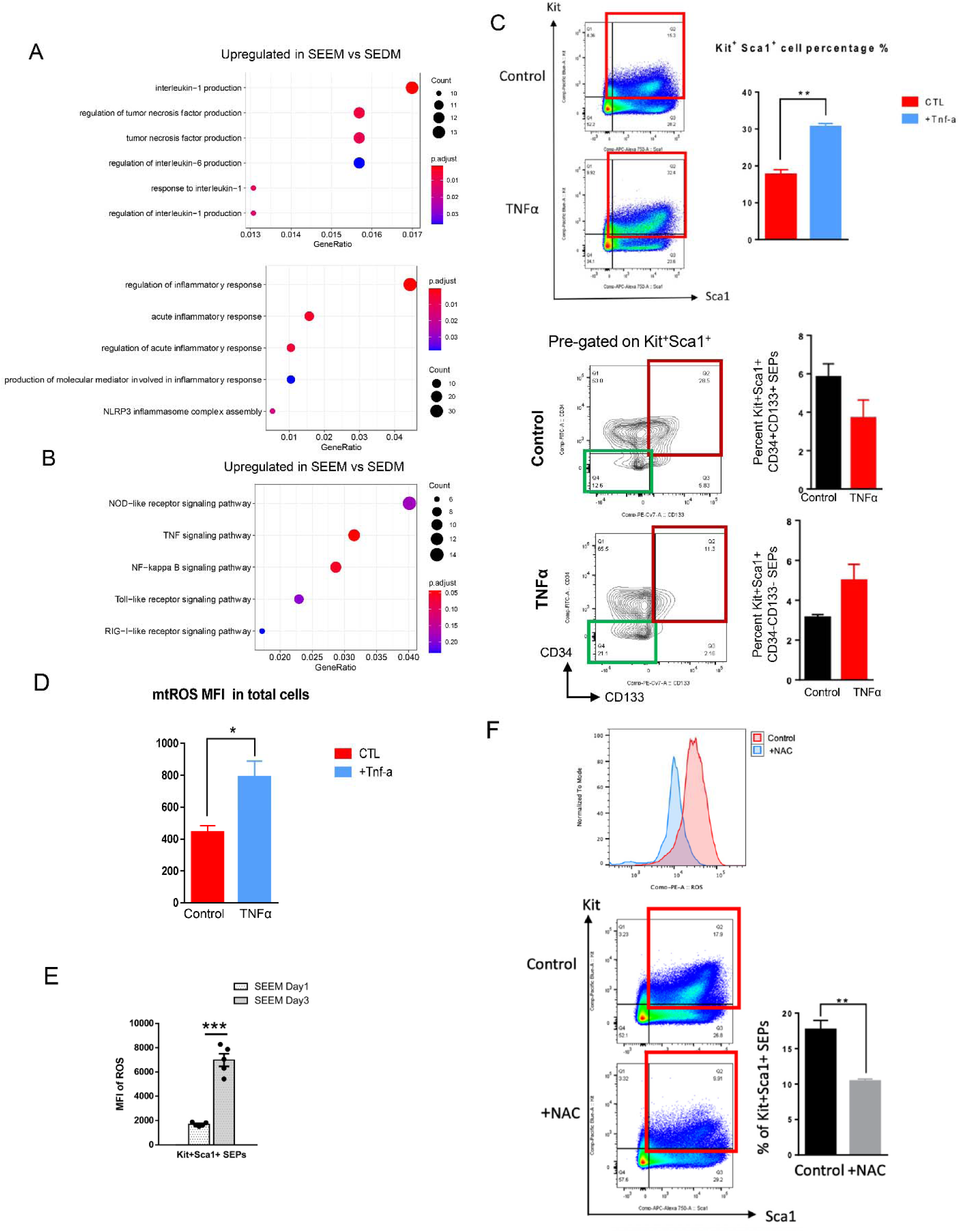
Immature SEPs utilize TNF-_α_-dependent production of ROS for SEP expansion and maturation. (A-B) RNA-seq analysis comparing SEPs isolated from SEEM and SEDM cultures. Overrepresentation analysis of upregulated DEGs in SEEM (FDR < 0.05, Fold change (FC) of SEEM/SEDM > 1.5), showing GO terms related to inflammatory response (A), and NF-κB- and PRRs-related KEGG pathways (B). Circle size represents the numbers of genes in each pathway and color represents BH-adjusted p value (n=3 per group). (C) SEPs were treated with ± 50ng/ml recombinant TNF-α protein at the start of SEEM culture for 48 hrs, after which SEPs were switched to fresh SEEM media for another 72 hrs. Flow cytometry analysis of Kit^+^Sca1^+^ SEPs (top) and analysis of pre-gated Kit^+^Sca1^+^cells with additional markers CD34 and CD133 (bottom) (n=3 per group, unpaired t test). (D) Quantification of mitochondrial ROS level by MFI of MitoSOX Red staining in SEPs treated with TNF-α protein as described in (C) (n=3 per group, unpaired t test). (E) SEPs isolated from SEEM cultures at day 1 and 3 were analyzed for intracellular ROS levels in Kit^+^Sca1^+^ SEPs. ROS levels were quantified by MFI of CellROX staining (n=5 per time point; unpaired t test). (F) SEPs were treated with ± 5 mM NAC at day 4 of SEEM culture for 24 hrs, followed by flow cytometry analysis of intracellular ROS levels (top) and analysis of percentages of Kit^+^Sca1^+^ SEPs (bottom) (n=3 per group, unpaired t test). Data represent mean ± SEM. * p < 0.05, ** p < 0.01, *** p < 0.001.

### Decreased itaconate production enables the elevated inflammatory signaling to drive SEP expansion

We hypothesized that the inflammatory response we observed in proliferating SEPs is modulated by intracellular metabolic changes. To this end, we performed LC-MS analysis to profile the changes of metabolites extracted from bulk SEPs on days 1 and 3 of SEEM cultures (see accompanying manuscript Ruan et al. submitted). We observed a striking decrease of an endogenous metabolite itaconate on day 3 of the expansion culture (Figure 2A).

**Figure 2.**
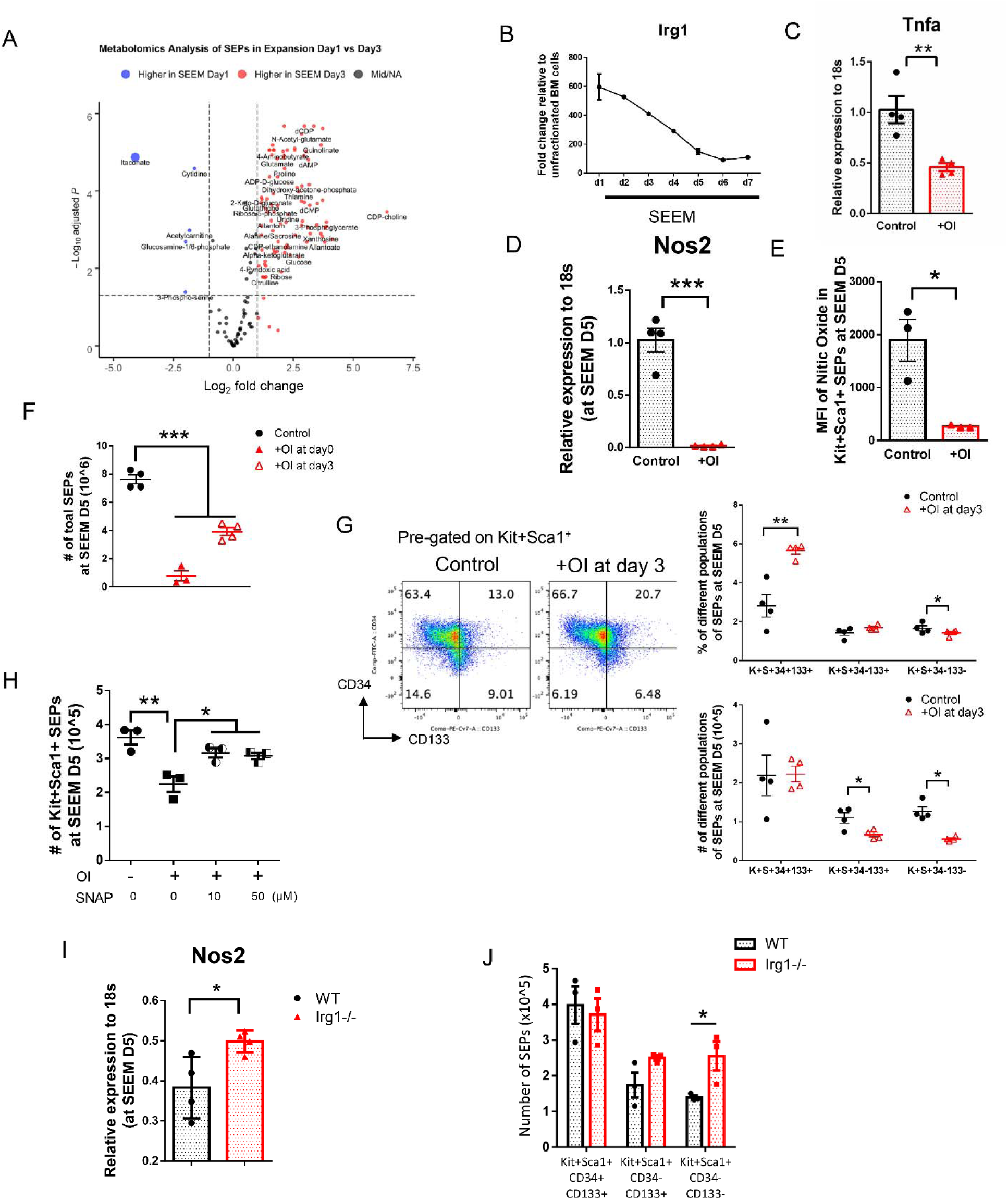
Decreased itaconate production enables the elevated inflammatory signaling to drive SEP expansion. (A) SEPs were isolated from SEEM cultures at day 1 and 3 for metabolomics analysis. Volcano plot showing the changes in metabolites between day 1 and 3 SEPs in SEEM (n=5 per time point). (B) qRT-PCR analysis of *Irg1* mRNA expression in SEPs from SEEM over time (n=3 per time point). (C-E) SEPs were treated ± 125 μM OI at SEEM day 3 for 48 hrs. Analysis of *Tnf-*_α_ (C) and *Nos2* (D) expression by qRT-PCR, and quantification of intracellular NO levels in Kit^+^Sca1^+^ SEPs by mean fluorescence intensity (MFI) of DAF-FM DA staining (E) (n=4 per group, unpaired t test). (F) SEPs were treated ± 125 μM OI at indicated days of SEEM cultures. On day 5 of SEEM cultures, total SEP cell counts were measured (n=4 per group, one-way ANOVA/Dunnett’s). (G) SEPs were treated ± 125 μM OI at SEEM day 3 for 48 hrs followed by flow cytometry analysis of SEPs. Representative flow cytometry plot showing pre-gated Kit^+^Sca1^+^cells with additional markers CD34 and CD133 (left), quantification of the percentages (top right) and absolute numbers (bottom right) of indicated populations of SEPs (n=4 per group, unpaired t test). (H) SEEM cultures were treated with 125 μM OI alone or in combination with SNAP for 24 hrs. Flow cytometry quantification of numbers of Kit^+^Sca1^+^ SEPs (n=3 per group, one-way ANOVA/Tukey’s). (I-J) WT and Irg1-/- SEPs were cultured in SEEM for 5 days followed by analysis Nos2 mRNA expression (I), and flow cytometry quantification of absolute numbers of indicated populations of SEPs (J) (n=4 per genotype, unpaired t test). Data represent mean ± SEM. * p < 0.05, ** p < 0.01, *** p < 0.001.

Immunoresponsive gene 1 (*Irg1*) encodes the enzyme that catalyzes itaconate synthesis from the decarboxylation of cis-aconitate, a TCA cycle intermediate(^29^). Correlated with a decline in itaconate, the mRNA expression of *Irg1* declined during the cultures in SEEM over time (Figure 2B).

In view of a well-established role of itaconate as an anti-inflammatory mediator(^30^), we reasoned that the decrease of itaconate production enables increased inflammatory signals to drive SEP proliferation. To test this possibility, we examined whether treating cultures with 4-octyl itaconate (OI), a cell-permeable itaconate derivative, decreases inflammatory signal-dependent SEP expansion. Our analysis showed that OI reduced the *Tnf-*_α_ mRNA expression in proliferating SEPs (Figure 2C). In addition to TNF-α, inducible nitric oxide synthase (*Nos2* or *iNOS*) derived nitric oxide (NO) is another key inflammatory signal necessary for expansion (see accompanying manuscript Ruan et al. submitted). Like *Tnf-*_α_, *Nos2* mRNA and NO levels were compromised by OI treatment (Figure 2D-E). OI-mediated suppression of TNF-α and NO in turn blocked SEP expansion (Figure 2F). In fact, if OI was added at the start of culture, SEPs failed to proliferate. In contrast to the effect of TNF-α, OI delayed the maturation of progenitors (Figure 1C and 2G). OI-treated SEEM cultures displayed fewer more rapidly proliferating late-stage Kit^+^Sca1^+^CD34^-^CD133^+^ and Kit^+^Sca1^+^CD34^-^CD133^-^ SEPs, while the numbers of the most immature Kit^+^Sca1^+^CD34^+^CD133^+^ progenitors were not affected (Figure 2G). Treatment with OI decreased the MFI of NO in SEPs (Figure S1B). In contrast, the defects in SEP expansion induced by OI treatment were rescued by treatment with the NO donor SNAP at either 10 or 50 μM, indicating that itaconate impairs proliferation by counteracting NO (Figure 2H). In contrast to OI treatment, culturing Irg1-/- cells resulted in increased expression of *Nos2* mRNA, increased SEP proliferation, and accelerated the transition to more mature Kit^+^Sca1^+^CD34^-^CD133^-^ cells (Figure 2I-J). Human stress erythropoiesis cultures exhibited a similar response to OI, which decreased proliferation of SEPs resulting in fewer late-stage progenitors (Figure S1C-D). Our findings reinforce the idea that pro-inflammatory signals promote the expansion and maturation of early SEP populations, while the anti-inflammatory metabolite itaconate acts to antagonize their effects.

### Itaconate impairs SEP expansion in a Nrf2-dependent manner

Itaconate alkylates Keap1 and acts as a potent inducer of Nrf2, which is a central regulator of the response to oxidative stress. Nrf2 is also involved in anti-inflammatory response, and this function is essential for the immunomodulatory role of itaconate in the activated macrophages(^31^). We hypothesized that itaconate inhibits SEP proliferation by driving the activation of Nrf2. Similar to what we observed with *Irg1* expression, mRNA expression of *Nrf2* and NAD(P)H:quinone oxidoreductase 1 (*Nqo1*), a Nrf2 target gene, decreased during cultures in SEEM (Figure 2B and 3A). Nrf2-deficient progenitors initially grew faster when compared to WT controls, however, there was no difference in total cell numbers at day 5 (Figure 3B). Nrf2-/- cultures contained more late-stage Kit^+^Sca1^+^CD34^-^CD133^-^ progenitors and fewer immature Kit^+^Sca1^+^CD34^+^CD133^+^ SEPs than WT cultures (Figure 3C). The similarities in expression and phenotypes of Irg1 and Nrf2 mutant progenitors suggests a model where low itaconate levels result in Nrf2 in an inactive state, which allows for SEP proliferation. To verify this mechanism, WT and Nrf2-/- SEEM cultures were supplemented with OI or dimethyl fumarate (DMF), a second known activator of Nrf2. Treatment of cultures with either OI or DMF led to increased *Nqo1* mRNA expression, which was blocked in Nrf2-/- SEPs (Figure 3D and S2A). Furthermore, the defect in SEP proliferation caused by addition of OI was rescued by mutation of Nrf2 in the Nrf2-/- SEPs (Figure 3E) and to a lesser extent in DMF treated cultures (Figure S2B). These data demonstrate that proliferation of SEPs during expansion phase relies on low Nrf2 activity.

**Figure 3.**
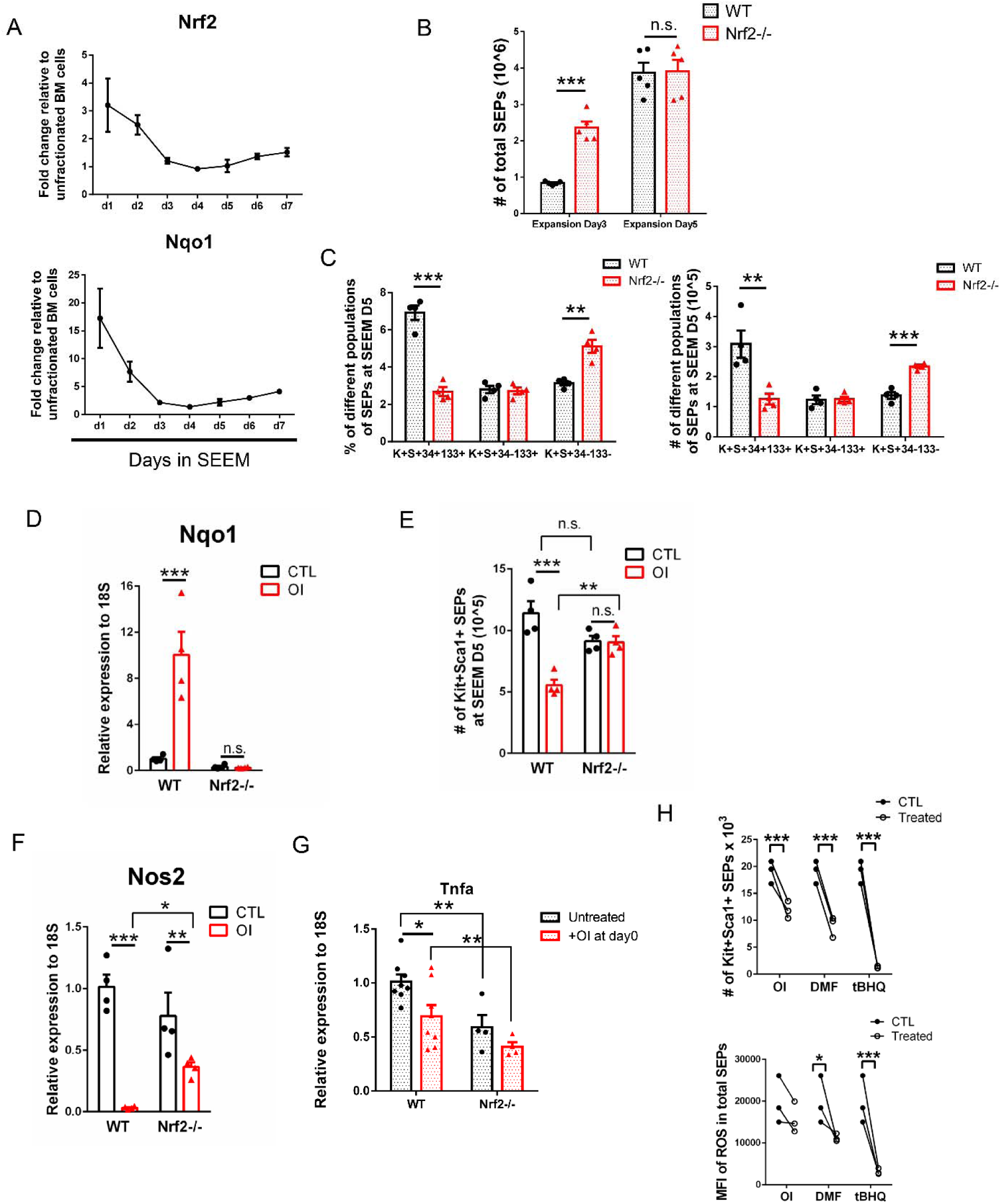
Itaconate impairs SEP expansion in a Nrf2-dependent manner. (A) qRT-PCR analysis of *Nrf2* (top) and *Nqo1* (bottom) expression in SEPs cultured in SEEM (n=3 per time point). (B) Analysis of total SEP counts in WT and Nrf2-/- SEEM cultures at day 3 and 5 (n=5 per group, unpaired t test). (C) Flow cytometry quantification of the percentages (left) and absolute numbers (right) of indicated populations of SEPs in WT and Nrf2-/- SEEM cultures at day 5 (n=4 per group, unpaired t test). (D-G) WT and Nrf2-/- SEEM cultures were treated ± 125 μM OI for 5 days. qRT-PCR analysis of *Nqo1* expression (D), analysis for numbers of Kit^+^Sca1^+^ SEPs (E), and qRT-PCR analysis of *Nos2* and *Tnf-*_α_ expression (n=4 (D-F), n=8 in WT and n=4 in Nrf2-/- (G); two-way ANOVA/Fisher’s LSD). (H) SEPs were treated with vehicle, 125 μM OI, 30 μM DMF or 20μM tBHQ at SEEM day 2 for 24 hrs, followed by flow cytometry analysis of numbers of Kit^+^Sca1^+^ SEPs (top) and MFI quantification of intracellular ROS levels (bottom) (n=3 per group, paired t test). Data represent mean ± SEM. n.s. p > 0.05, * p < 0.05, ** p < 0.01, *** p < 0.001.

We next examined whether Nrf2 suppresses the inflammatory signals required for SEP expansion. As we observed with OI, DMF treatment also decreased NO levels in progenitor cells (Figure S2C). In addition, either OI or DMF treatment decreased expression of *Nos2* mRNA, and this effect was compromised in the Nrf2-/- SEPs (Figure 3F and S2D). These results demonstrate that active Nrf2 inhibits iNOS-derived NO production. Like the decrease in *Nos2* expression induced by OI, the decrease in *Tnf-*_α_ mRNA expression induced by OI was reversed by Nrf2 mutation (Figure 3G). In contrast to OI, DMF treatment increased *Tnf-*_α_ expression (Figure S2E). Although OI and DMF have different effects on *Tnf-*α expression, treating SEEM cultures with OI, DMF or another Nrf2 activator tert-Butylhydroquinone (tBHQ) decreased the proliferation of Kit+Sca1+ SEPs and blocked the accumulation of intracellular ROS (Figure 3H). Our previous data showed that in immature SEPs mRNA expression of *Hif-1*_α_ and *Pdk1* promotes glycolysis, which provides anabolic metabolites for cell proliferation(^16^). OI or DMF treatment decreased *Hif-1*_α_ and *Pdk1* mRNA expression in proliferating SEPs and this decrease was reversed by Nrf2 mutation (Figure S3A-B), suggesting that activation of Nrf2 disrupts the inflammatory metabolism required for SEP proliferation.

Collectively, these data support a model where low levels of itaconate maintain Nrf2 in an inactive state, which allows for NO- and ROS-dependent proliferation of SEPs during the initial expansion stage of stress erythropoiesis.

### Epo initiates an itaconate-dependent anti-inflammatory response to promote SEP differentiation

Previously we showed that Epo signaling promotes the transition of proliferating progenitors to erythroid differentiation(^18, 19, 32^). Switching stress erythropoiesis cultures to differentiation media led to an increase of erythroid gene expression (Figure 4A), whereas it decreased the expression of pro-inflammatory genes (Figure 1A-B). The resolution of inflammation was coupled to a profound switch of metabolism, in which the level of itaconate increased after switching to SEDM (Figure S4A). Consistently, the mRNA expression of *Irg1* also increased (Figure 4B). These data suggest that the Epo induced transition to differentiation increases the production of itaconate that drives an anti-inflammatory response to promote SEP differentiation. We performed SEDM cultures using control or Irg1-/- bone marrow cells in which itaconate production was completely impaired. We restored the levels of itaconate in Irg1-/- cultures with the supplementation of OI. Compared to controls, Irg1-/- SEPs had elevated levels of iNOS protein and NO production, but treatment with OI decreased iNOS protein and NO levels to levels comparable to control cells (Figure 4C-D and S4B). To test whether itaconate promotes differentiation via NO suppression, we isolated SEPs from control and Irg1-/- SEDM cultures treated with and without iNOS inhibitor 1400w. Irg1-deficient progenitors generated fewer stress BFU-Es and mature Kit^+^Sca1^-^CD34^-^CD133^-^ SEPs and expressed lower mRNA levels of erythroid differentiation-associated genes, indicating that itaconate production is required for the transition to erythroid differentiation (Figure 4E-G). In contrast, treatment with 1400w, a Nos2 specific inhibitor, rescued the defects in erythroid differentiation of Irg1-deficient SEPs. Similar to iNOS protein, ROS levels also decreased following SEP differentiation (Figure S4C).

**Figure 4.**
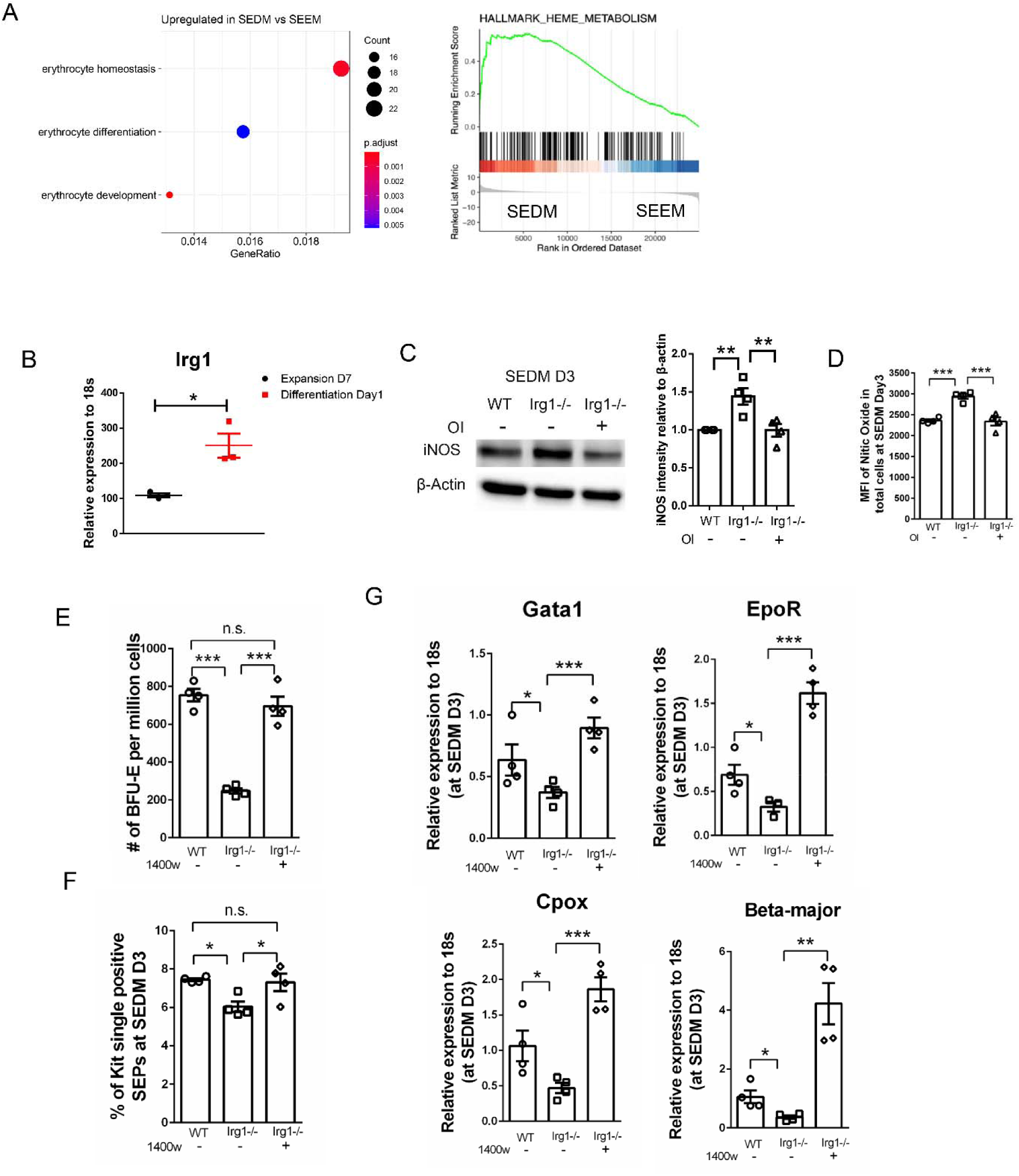
Increased itaconate production during differentiation alleviates NO-dependent erythroid inhibition. (A) RNA-seq analysis comparing SEPs from SEEM and SEDM cultures. Overrepresentation analysis of upregulated DEGs in SEDM (FDR < 0.05, FC of SEDM/SEEM > 1.5), showing significant enriched erythroid-associated GO terms. Circle size represents the numbers of genes in each GO term and color represents adjusted p value (left). Molecular Signatures Database (MSigDB)-based GSEA analysis showing heme metabolism (middle) (n=3 per group). (B) qRT-PCR detection of progenitor *Irg1* expression between SEEM Day7 and SEDM Day1 (n=3 per group, unpaired t test). (C-D) SEPs were harvested from WT, or Irg1-/- SEDM cultures treated ± 125 μM OI for 3 days. WB (left) and densitometry band quantification (middle) of iNOS protein expression (C), and quantification of NO levels (D) (n=4 per group, one-way ANOVA/Tukey’s). (E-G) SEPs were isolated from WT, or Irg1-/- SEDM cultures treated ± 10 μM 1400w for 3 days. Analysis of stress BFU-E by colony assay (E), flow cytometry analysis of the percentages of Kit^+^Sca1^-^CD34^-^CD133^-^ SEPs (F) and qRT-PCR detection of erythroid-specific genes (G) (n=4 per group, one-way ANOVA/Tukey’s). Data represent mean ± SEM. n.s. p > 0.05, * p < 0.05, ** p < 0.01, *** p < 0.001.

However, unlike decreasing NO which leads to increased differentiation, decreasing ROS by treating proliferating SEPs with NAC 24 hours prior to the switch to differentiation media, significantly reduces differentiation as demonstrated by fewer stress BFU-E and reduced erythroid gene expression (Figure S4D-F). These data demonstrate that itaconate promotes erythroid differentiation by inhibiting iNOS-dependent NO production, but the role of ROS in regulating differentiation is more complex.

We next investigated the role of Irg1 *in vivo* during the recovery from Heat Killed *Brucella abortus* (HKBA) induced inflammatory anemia(^33–35^). Untreated Irg1-/- mice exhibited similar levels of SEPs in their spleens and stress BFU-E when compared to wildtype (Figure S4G). Despite this similarity, Irg1-/- mice treated with HKBA exhibited a significant delay in recovery over 28 days (Figure 5A). On day 8 after HKBA treatment, the anemia of control mice starts to improve, and over the next 8 days the mice significantly improve their hematocrit. We examined Irg1-/- and control HKBA treated mice during this critical period in recovery on days 8, 12 and 16. We observed that Irg1-/- showed continued decreases in hemoglobin and RBC concentration during this time (Figure 5B). However, this defect of stress erythropoiesis is not due to a lack of SEPs in the spleen as spleen weight and spleen cellularity was increased in the Irg1-/- mice (Figure 5C). The defect is in the differentiation of SEPs. The percentage of Kit^+^Sca1^-^CD34^-^ CD133^-^ SEPs was significantly decreased in Irg1-/- mice, while the percentage of Kit+Sca1+CD34^+/-^ CD133+ immature cells increased (Figure 5D-E). This decrease in mature SEPs translated to a lower frequency of stress BFU-E at each time point and fewer overall stress BFU-E on days 8 and 16 (Figure 5F). Analysis of *Nos2* expression in the spleens showed that Irg1-/- mice had increased levels of Nos2 supporting the role for itaconate synthesis in suppressing NO dependent inhibition of erythroid differentiation (Figure 5G).

**Figure 5.**
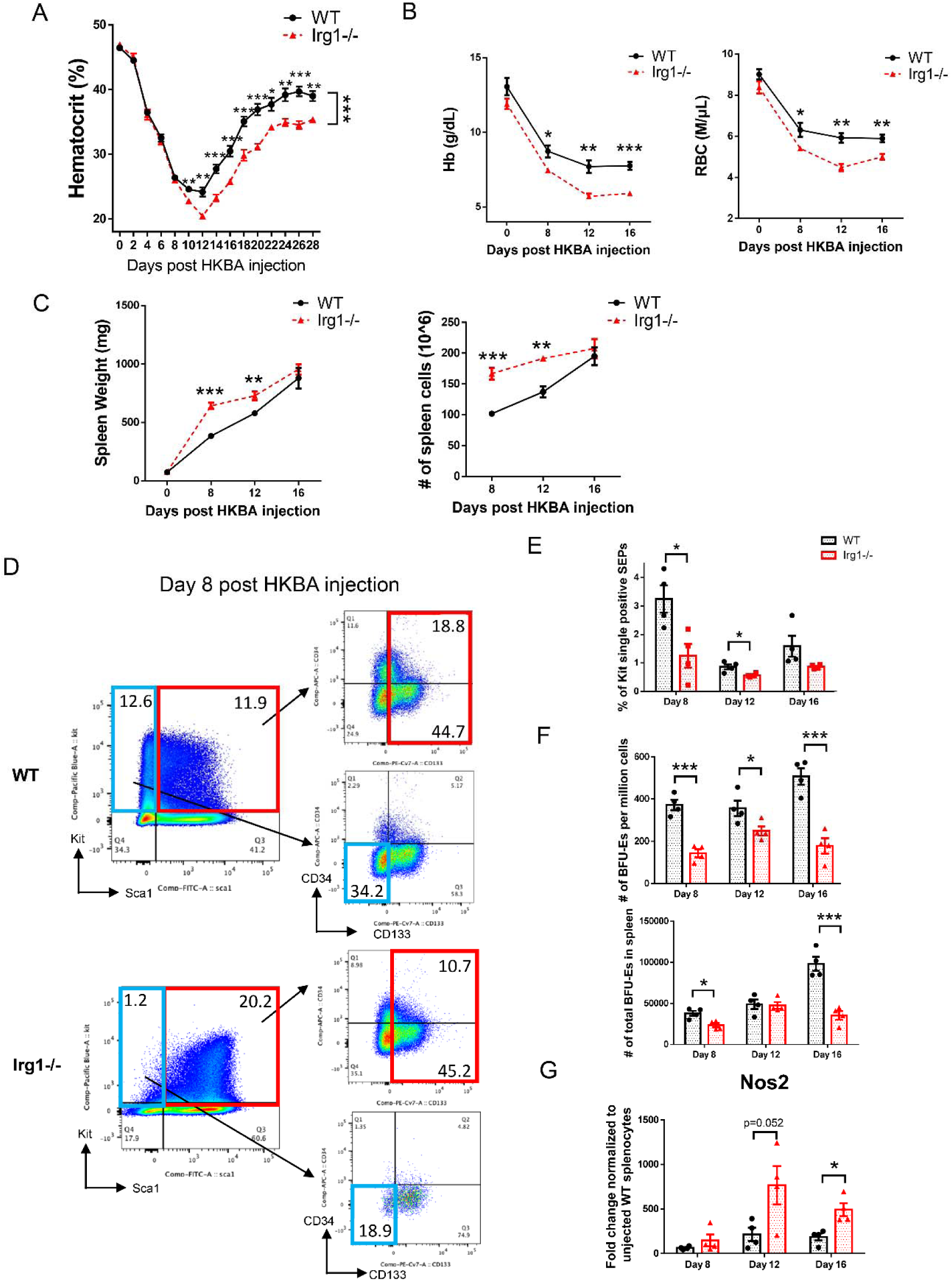
Defective SEP differentiation in Irg1-deficient mice delayed the recovery from HKBA-induced inflammatory anemia. (A-G) Age- and sex-matched WT and Irg1-/- mice were administered with HKBA (5 x 10^8^ particles/mouse) via intraperitoneal injection. (A) In the following 28 days, mice were monitored daily for survival and health, and blood was collected retro-orbitally in every other day for measurement of hematocrit (n=12 in WT, n=8 in Irg1-/-, repeated measures two-way ANOVA followed by unpaired t test). (B-C) Analysis of Hb (left) and RBC counts (right) concentrations (B), and measurement of spleen weight (left) and splenocyte numbers (right) (C) at indicated time points post HKBA injection (n=4 per group, unpaired t test). (D) Representative flow cytometry plot showing the gating of SEPs in the spleen isolated at day 8 post HKBA injection. (E-G) Analysis of percentages of Kit^+^Sca1^-^CD34^-^CD133^-^ SEPs (E), frequency (top) and total numbers (bottom) of stress BFU-E colony formation (F), and *Nos2* mRNA abundance (G) at indicated time points (n=4 per group, unpaired t test). Data represent mean ± SEM. * p < 0.05, ** p < 0.01, *** p < 0.001.

### Itaconate and IL-10 work in concert to activate Nrf2-mediated SEP differentiation

RNA-seq analysis comparing gene expression of Nrf2-/- and control SEPs isolated from SEDM cultures showed that Nrf2-deficient progenitors were significantly enriched in pathways associated with inflammatory response, as well as NF-κB and pro-inflammatory cytokine-related signaling (Figure 6A and S5A-B). In contrast, genes associated with erythroid differentiation and hemoglobin biosynthesis were severely decreased by Nrf2 deficiency (Figure 6B). Furthermore, Nrf2-/- SEPs expressed lower levels of genes involved in ribosome biogenesis and amino acid metabolism, suggesting a decline of translational efficiency for hemoglobin production (Figure S5B-D). The RNA-seq data indicate that Nrf2 promotes an anti-inflammatory response to promote SEP differentiation. To demonstrate that itaconate dependent activation of Nrf2 drives differentiation, we isolated SEPs from control and Irg1-/- SEDM cultures supplemented with OI or DMF. Compared to controls, Irg1-/- SEPs expressed lower mRNA levels of Nrf2 targets, which were induced by treatment with exogenous OI and DMF (Figure S6A). Furthermore, we observed that the treatment with OI and to a lesser extent DMF of Irg1-deficient progenitors rescued the defects in erythroid gene expression and stress BFU-Es (Figure 6C-D and S6B). These data demonstrate that the role of itaconate in promoting differentiation is Nrf2 dependent.

**Figure 6.**
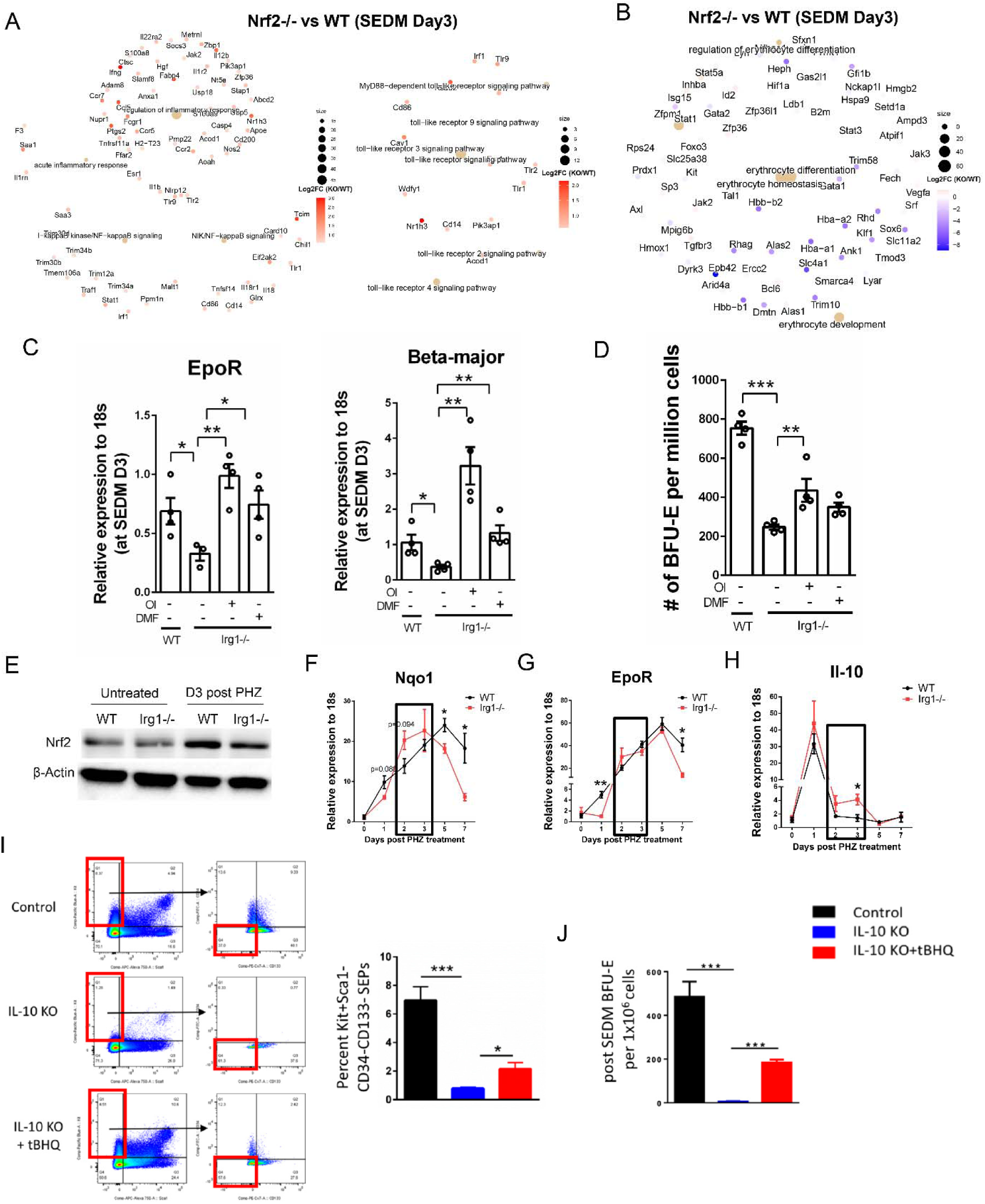
Anti-inflammatory signals itaconate and IL-10 act in concert to activate Nrf2-mediated SEP differentiation. (A-B) RNA-seq analysis comparing WT and Nrf2-/- SEPs isolated from SEDM cultures. Network analysis of upregulated DEGs in Nrf2-/- (FDR < 0.05, FC of Nrf2-/-/WT > 1.5) in pathways associated with inflammatory response and toll-like receptor signaling (A). Network analysis of DEGs (FDR < 0.05, |FC| > 1.2) in erythroid-associated pathways (B). Color representing log2FC of SEDM Nrf2-/-/WT (n=3 per genotype). (C-D) SEPs were harvested from WT, or Irg1-/- SEDM cultures treated with vehicle, 125 μM OI or 30 μM DMF for 3 days. Analysis of mRNA expression of erythroid-specific genes by qRT-PCR (C). Analysis of stress BFU-E by colony assay (D) (n=4 per group, one-way ANOVA/Tukey’s). (E-H) Age- and sex-matched WT and Irg1-/- mice were injected intraperitoneally with a single dose (100 mg/kg body weight) of freshly prepared phenylhydrazine. WB analysis of Nrf2 protein levels in day 0 (untreated) and day 3 post PHZ injection (E). qRT-PCR detection of mRNA levels of *Nqo1* (F), *EpoR* (G) and *Il-10* (H) (n=3 per time point, unpaired t test). (I-J) SEPs were isolated from WT, or IL-10-/- SEDM cultures treated ± 20μM tBHQ for 3 days. Representative flow cytometry plot (left) and quantification of Kit^+^Sca1^-^CD34^-^CD133^-^ SEPs (right) (I), and the frequency of stress BFU-E colony formation (J) (n=4 per group, unpaired t test). Data represent mean ± SEM. * p < 0.05, ** p < 0.01, *** p < 0.001.

Nrf2 protein levels increased in the spleen during the recovery from PHZ-induced anemia (Figure 6E and S6C). Consistently, we observed a robust induction of Nrf2 target genes *in vivo* (Figure 6F and S6D-E). In contrast, Irg1 mutants failed to induce Nrf2 protein to comparable levels as WT controls (Figure 6E). The mRNA expression of Nrf2 responsive genes also decreased in Irg1-/- mice, however this decrease occurred only in the early time points and expression increased at later time points during recovery (Figure 6F). We also observed that the defects in erythroid differentiation were transient and the Irg1-/- mice eventually reached levels of erythroid gene expression, stress BFU-E and hematocrit similar to controls (Figure 6G and S6F-G). These data suggest that an alternative mechanism to activate Nrf2 compensates during the recovery from acute anemia. This observation is also supported by *in vitro* data where we observed that the defects in erythroid gene expression in Irg1-/- SEPs are not as severe as Nrf2-/- cells (Figure 6B-C). In addition to itaconate, the expression of another anti-inflammatory signal, IL-10, increased when stress erythropoiesis cultures were switched to SEDM (Figure S6H). Irg1-/- mice had higher *Il-10* mRNA levels than control mice between day 2 and day3 post PHZ injection, and this period is the time we observed the compensation of Nrf2 activity (Figure 6F-H). To test whether IL-10 is an alternative signal to activate Nrf2, we harvested SEPs from control and IL-10-/- SEDM cultures treated with and without Nrf2 activator tBHQ. IL-10-/- SEPs had lower mRNA expression of Nrf2 targets when compared to control cells, but it was restored by tBHQ treatment (Figure S7A). In addition, IL-10 deficiency resulted in fewer mature Kit^+^Sca1^-^ CD34^-^CD133^-^ SEPs when cells were grown in differentiation media, which translated into fewer stress BFU-E. Lack of IL-10 also resulted in decreased expression of erythroid-specific genes (Figure 6I-J and S7A). However, treatment with tBHQ to activate Nrf2 in IL-10-/- cultures rescued the defects in erythroid differentiation. Treatment with an inhibitor for the IL-10 receptor to disrupt IL-10 signaling reproduced the defects of IL-10-/- SEPs in terms of mRNA expression of Nrf2 targets and erythroid genes (Figure S7B). Collectively, these findings demonstrate that itaconate and IL-10 act in concert to activate Nrf2, which drives an anti-inflammatory response to promote the transition of progenitors to differentiate.

### Nrf2-dependent regulation of SEP differentiation acts on both progenitors and niche cells

The development of SEPs requires a close interaction with the macrophages in the stress erythropoiesis niche(^14^). Previously, work from Gbotosho et al. showed a requirement for Nrf2 signaling in macrophages during stress erythropoiesis(^36^). To determine whether Nrf2 acts on SEPs or niche cells, we performed co-culture experiments in which CD45.1 WT or CD45.2 Nrf2-/- stromal cells were seeded with progenitor cells from the same or opposite genetic background. Our co-culture experiment showed that Nrf2 deficiency in SEPs alone decreased the production of Kit^+^Sca1^-^CD34^-^CD133^-^ cells, BFU-E and expression of erythroid markers (Figure 7A-C and S7C), indicating a progenitor cell-intrinsic role of Nrf2. Additionally, Nrf2 deficiency in both SEPs and stromal cells led to the strongest defects in erythroid differentiation, which is consistent Gbotosho’s work suggesting that niche macrophages require Nrf2 signaling to regulate stress erythropoiesis(^36^).

**Figure 7.**
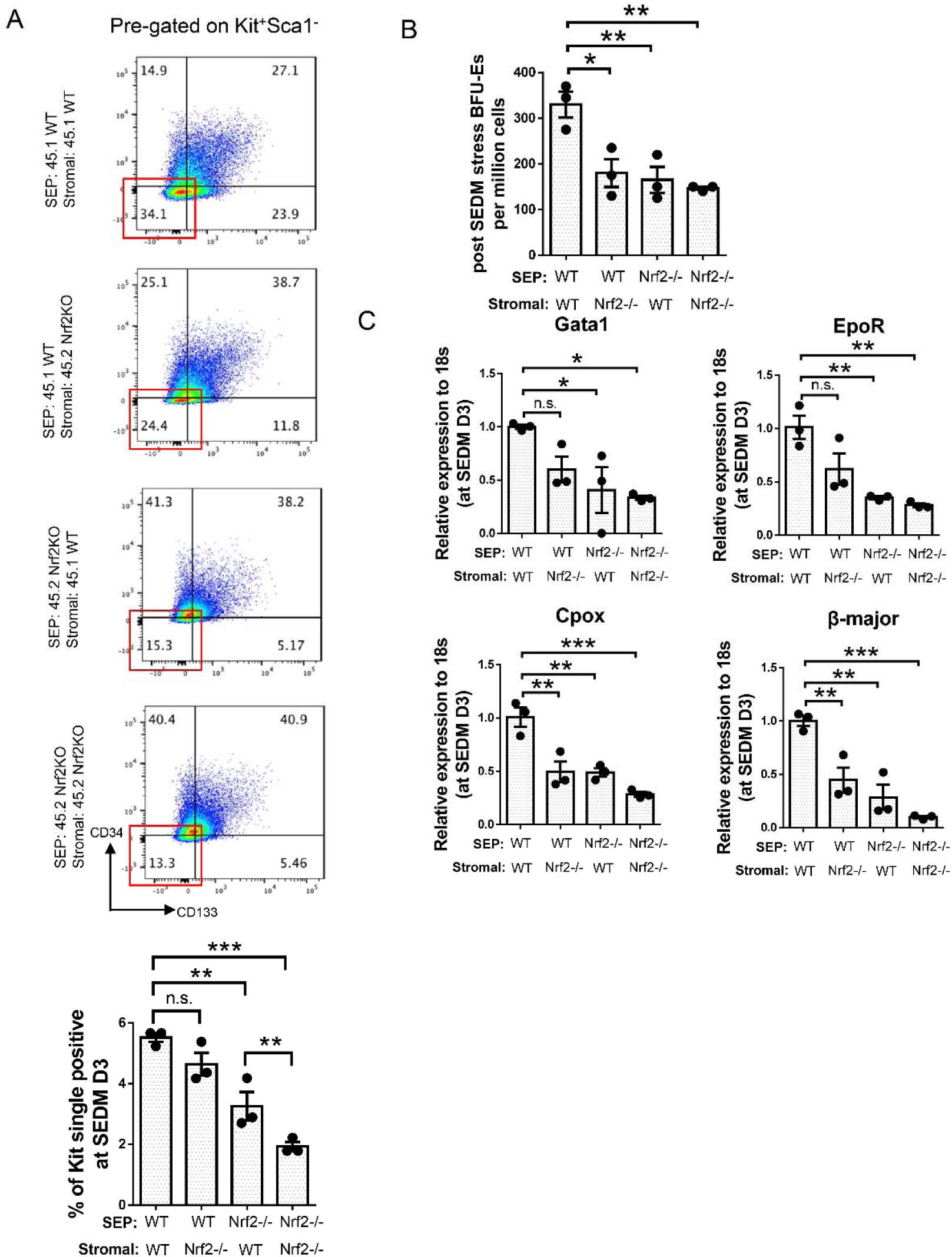
Nrf2-dependent regulation of SEP differentiation acts on both progenitors and niche cells. (A-C) WT (CD45.1) and Nrf2-/- (CD45.2) BM cells were cultured in SEEM for 5 days followed by 3 days in SEDM. When switched to SEDM, nonadherent SEPs were collected and plated on the stromal layer from indicated genotypes. (A) SEPs were isolated from SEDM cultures for flow cytometry analysis. CD45.1 was stained to gate SEPs that were derived from the seeded non-adherent cells (Nrf2-/- CD45.2; WT CD45.1), followed by gating on Kit and Sca1. Pre-gated Kit+Sca1-cells were next gated on CD34 and CD133. Representative flow cytometry plot (top) and quantification of the percentages of Kit+Sca1-CD34-CD133-SEPs (bottom) (n=3 per group, one-way ANOVA/Tukey’s). (B) Analysis of the frequency of stress BFU-E colony formation (n=3 per group, one-way ANOVA/Tukey’s). (C) qRT-PCR analysis of mRNA expression of indicated erythroid-specific genes (n=3 per group, one-way ANOVA/Tukey’s). Data represent mean ± SEM. n.s. p > 0.05, * p < 0.05, ** p < 0.01, *** p < 0.001.

## Discussion

Inflammation enhances the production of inflammatory cytokines and chemokines from red-pulp macrophages, which in turn mobilizes an influx of bone marrow monocytes into the spleen to build on the early inflammatory niche(^14, 15^). In this study, we show that early SEPs respond to niche signals by increasing the expression of pro-inflammatory genes, Tnf-α and Nos2, to boost the production of ROS and NO respectively. NO and ROS serve as signaling molecules to drive the proliferation of early SEPs. NO also inhibits erythroid differentiation (See companion manuscript Ruan et al.). Consequently, inflammation expands a large pool of immature SEPs prior to their differentiation. Failure to induce inflammatory signals in early SEPs, due to treatment of anti-inflammatory molecule OI or DMF, blocks SEP proliferation, which in vivo would result in too few erythrocytes being produced to restore homeostasis. Our data show that inflammation induces proliferation of SEPs but inhibits their differentiation, which is consistent with murine studies using different infection and sterile inflammation models, where pro-inflammatory cytokines increase myelopoiesis, but suppress bone marrow erythropoiesis. To compensate for the loss of erythroid output, pro-inflammatory signals drive the expansion of immature SEP populations resulting in extramedullary stress erythropoiesis(^15, 33–35, 37^). Together, these results highlight stress erythropoiesis as a major component of inflammatory response, which functions to balance the physiological needs of erythrocytes for tissue oxygenation with the need to produce myeloid effector cells to fight infection and heal wounds.

Here we show that the transition from proliferating SEPs to committed progenitors requires a switch in signals from pro-inflammatory to pro-resolving signals. This transition response is mediated in part by the immunomodulatory molecule itaconate. Itaconate is a well-known anti-inflammatory metabolite that affects the activities of a number of immune cells including macrophages and dendritic cells(^23^). Our data here showed that itaconate, which is known to alkylate Keap1 preventing its interaction with Nrf2(^31^), activates Nrf2 leading to the repression of *Nos2* expression. However, itaconate has many more targets and can affect metabolism by directly inhibiting succinate dehydrogenase(^38^). Recent work from Marcero et al. showed that itaconate can also inhibit erythroid differentiation by blocking heme biosynthesis through the inhibition of ALAS2(^39^). In that study, levels of TCA metabolites were altered in erythroid cultures treated with itaconate. These data suggest that itaconate synthesis must be carefully regulated, although it is required for the transition from NO-dependent proliferation of SEPs to the Epo-dependent commitment to erythroid differentiation, too much itaconate could compromise heme biosynthesis and inhibit terminal differentiation. Our data focuses on the role of itaconate in SEPs, but itaconate could also affect the niche. Itaconate decreases the output of pro-inflammatory cytokines by M1 macrophages(^23, 30^). Our previous work showed that the transition to differentiation relies on a change in the niche from one dominated by pro-inflammatory signals to one dominated by anti-inflammatory signals(^15, 19^). Activation of PPARγ by prostaglandin J2 is a key signal in this change(^19^). Mutation of PPARγ leads to increased *Irg1* expression and higher levels of Itaconate(^40^). The switch from M1 to M2 like macrophages in the splenic stress erythropoiesis niche may rely on decreasing itaconate levels, which would also prevent the effects of prolonged itaconate signaling on erythroid differentiation.

In summary, our data show that inflammatory signals that induce NO and ROS production are utilized by stress erythropoiesis to expand a bolus of immature progenitors, whereas a tightly regulated resolution of inflammation is needed to ensure a successful transition of early progenitors into mature erythrocytes to restore homeostasis.

## Acknowledgments

We thank the members of the Paulson lab for suggestions on the work, especially Yuting Bai for her help with mice blood collection. We thank Sougat Misra for helpful discussion about metabolism, Philip Smith and Justin Munro for running LC-MS. This work was supported by NIH grant HL146528 (RFP), NIFA-USDA Hatch Project PEN04771 accession #0000005 (RFP and KSP), R01CA239256 (MH), NIFA-USDA Hatch Project PEN04275 accession #1018544 (MH), startup funds from the College of Agricultural Sciences, Pennsylvania State University (MH), the Dr. Frances Keesler Graham Early Career Professorship from the Social Science Research Institute, Pennsylvania State University (MH), the NIFA-USDA Hatch Project PEN04607 accession number 1009993 (ADP) and from the PA Department of Health using Tobacco CURE funds (ADP, IK).

## Authorship

B.R., K.S.P and R.F.P. conceived and designed the study. B.R. and Y.C. performed the experiments. B.R., J.M., and M.A.H. analyzed the RNA-seq data. B.R., I.K., J.C., and A.D.P. analyzed the metabolomics analysis. B.R. and R.F.P. wrote the initial draft of the manuscript, and all authors were involved in review and editing.

Conflict-of-interest disclosure: The authors declare no competing financial interests.

