## Supplemental methods and data for "Itaconate promotes the differentiation of murine stress erythroid progenitors by increasing Nrf2 activity"

Ruan et al.

Supplemental methods and data

### Supplemental Methods

#### Stress erythropoiesis expansion and differentiation medium[(1)]

Stress erythropoiesis expansion medium (SEEM) was prepared by supplementing IMDM with 10% (v/v) FBS, 0.0007% (v/v) 2-mercaptoethanol, 0.01 g/ml BSA, 10 μg/ml ciprofloxacin, 2 mM L-glutamine, 10 μg/ml insulin, 200 μg/ml holo-transferrin, 50 ng/ml murine SCF, 15 ng/ml human BMP4, 25 ng/ml murine SHH and 30 ng/ml murine GDF15. Stress erythropoiesis differentiation medium (SEDM) was prepared by additionally supplementing SEEM with 3 U/ml human Epo. SEEM were cultured in ambient air (20% $O_2$), while SEDM were cultured in a hypoxia chamber (2% $O_2$) to maximize differentiation potential. All cultures were incubated in 5% $CO_2$ at 37 °C.

Human SEEM was prepared by supplementing IMDM with 10% (v/v) embryonic stem-cell FBS, 0.0007% (v/v) 2-mercaptoethanol, 20% (v/v) BIT 9500 serum substitute, 10 μg/ml ciprofloxacin, 2 mM L-glutamine, and human recombinant proteins including 50 ng/ml SCF, 15 ng/ml BMP4, 25 ng/ml SHH and 30 ng/ml GDF15.

#### Stress erythropoiesis cultures

Bone marrow cells were isolated, and cells were plated into stress erythropoiesis expansion media (SEEM) at a starting concentration of 6 x $10^5$ cells/ml for a 5-day culture. For differentiation cultures, non-adherent cells harvested from SEEM cultures were resuspended in stress erythropoiesis differentiation media (SEDM) at 3 x $10^5$ cells/ml for another 3 days. Primary human bone marrow mononuclear cells (BMNCs) were used for human stress erythropoiesis cultures. BMNCs were adjusted to a concentration of 1 x $10^6$ cells/ml in human SEEM for 7 days. SEEM were cultured in ambient air (20% $O_2$), while SEDM were cultured in a hypoxia chamber (2% $O_2$). All cultures were incubated in 5% $CO_2$ at 37 °C.

#### In vivo induction of stress erythropoiesis

Phenylhydrazine was used to induce stress erythropoiesis in the context of acute hemolytic anemia. Mice were injected intraperitoneally with a single dose (100 mg/kg body weight) of freshly prepared phenylhydrazine (Sigma-Aldrich, dissolved in PBS)[2]. Blood and spleen samples were collected at indicated time points post injection for downstream analysis.

Heat-killed Brucella abortus (HKBA, strain 1119-3) was used to induce anemia of inflammation according to a previously described method[3]. After centrifuging, HKBA was resuspended in PBS to make a stock solution with a concentration of 5 x $10^9$ particles/ml. HKBA stock was 1:1 diluted with PBS before use. To induce stress erythropoiesis, age- and sex-matched mice were administered with 200 μl diluted HKBA (5 x $10^8$ particles/mouse) via intraperitoneal injection. In the following 28 days, mice were monitored daily for survival and health, and blood was collected retro-orbitally in every other day for microhematocrit test. To assess stress erythropoiesis, mice were sacrificed at indicated time points for blood and spleen collection.

**Microhematocrit centrifuges and complete blood count test**

To determine hematocrit levels by microhematocrit method, the peripheral blood was collected retro-orbitally with the heparin-coated microhematocrit tube (VWR). The tube with one end sealed (CRITOSEAL, Leica Microsystems) was centrifuged for 5 min at 11,700 rpm (Autocrit Ultra 3 microhematocrit centrifuge, BD Biosciences) to separate the blood sample. The hematocrit level was quantified as the percentage of the volume of packed red blood cells relative to the volume of whole blood. For complete blood count analysis, mouse blood was collected retro-orbitally with a $K_2$EDTA-coated Microtainer tube, and sample was immediately analyzed on a Hemavet 950 analyzer (Drew Scientific).

**Stress BFU-E colony assay[2]**

SEPs isolated from SEDM cultures or mouse splenocytes were counted using a hemocytometer. For each sample, 2.5 x $10^5$ cells were resuspended in 2 ml MethoCult M3334 media (STEMCELL Technologies) supplemented additionally with 50 ng/ml SCF (GoldBio) and 15 ng/ml BMP4 (Thermo Fisher Scientific), and cell suspension was evenly plated into 3 wells of a 12-well plate as technical triplicates. Stress BFU-Es were stained with benzidine and quantified after a 5-day culture in 37 °C with 2% $O_2$ and 5% $CO_2$.

**Flow cytometry analysis**

Cells were stained with Zombie Yellow Fixable Viability Kit (BioLegend) to exclude dead cells. After 15 min incubation at room temperature in the dark, cells were washed and prepared in single cell suspension in flow cytometry staining buffer. The combinations of fluorophore-conjugated cell surface antibodies were added to the cell suspension. For analysis of mouse stress erythroid progenitors, the following antibodies were used: Kit Brilliant Violet 421 (Clone 2B8, BioLegend), Sca-1 APC/Cyanine7 (Clone D7, BioLegend), Sca-1 FITC (Clone D7, BioLegend), CD34 Alexa Fluor 647 (Clone RAM34, BD Biosciences), CD34 FITC (Clone RAM34, BD Biosciences), and CD133 PE/Cyanine7 (Clone 315-2C11, BioLegend). For CD45.1 WT/CD45.2 Nrf2-/-co-culture experiment, the above antibodies were added together with CD45.2 FITC (Clone 104, BD Biosciences) to determine the cell sources. For analysis of human progenitors, c-Kit Brilliant Violet 421 (Clone 104D2, BioLegend), CD34 Alexa Fluor 647 (Clone 561, BioLegend) and CD133 PE (Clone 7, Clone 7 BioLegend) were used in combination. After 30 min incubation on ice in the dark, cells were washed twice and resuspended in 250 μl staining buffer. Flow cytometry analysis was performed on a BD LSR Fortessa Cytometer (BD Biosciences) and data were analyzed by FlowJo software (BD Biosciences). Intracellular NO, cellular ROS, and mitochondrial ROS levels were analyzed by flow cytometry using the fluorescent probe DAF-FM diacetate, CellROXGreen Flow Cytometry Assay Kit, and MitoSOX Red Mitochondrial Superoxide Indicator, respectively, according to vendor's instruction (Thermo Fisher Scientific).

**qRT-PCR**

Total RNA was extracted with TRIzol reagent (Thermo Fisher Scientific). 1 μg of total RNA was reverse transcribed to cDNA using the qScript cDNA Synthesis Kit (Quanta Biosciences). TaqMan Gene Expression assays were performed on a StepOnePlus Real-Time PCR System

(Applied Biosystems) using the PerfeCTa qPCR SuperMix ROX (Quanta Biosciences). Relative gene expression was quantified by the $\Delta\Delta C_T$ method in reference to the housekeeping gene 18S rRNA for normalization. See Table 2-2 for TaqMan probes.

**Western blot**

Spleen cells or cultured progenitor cells were lysed with RIPA Buffer (Thermo Fisher Scientific) in combination with protease inhibitor cocktail (Sigma-Aldrich) and PMSF (Cell Signaling Technology). Cultured cell lysates were vortexed and spleen cell lysates were briefly sonicated using a Bioruptor Standard Sonicator. Samples were incubated on ice for 30 min and centrifuged at 13,000 g for 15 min at 4°C. Protein concentration was quantified using the Pierce BCA Protein Assay Kit (Thermo Fisher Scientific) following manufacturer's protocol. 30 µg of total proteins were subjected to SDS-PAGE and transferred onto a PVDF membrane. The membranes were blocked with 5% milk in TBST at room temperature for 1 hrs and immunoblotted with primary antibodies against Nrf2 (1:1000, Proteintech), iNOS (1:1000, Cayman Chemical) and β-Actin (1:2000, Santa Cruz) overnight at 4°C. The blots were washed with TBST for three times, followed by incubation with HRP-conjugated secondary antibodies (1:5000, Thermo Fisher Scientific) for 1 hrs at room temperature. The blots were developed using SuperSignal West Pico PLUS Chemiluminescent Substrate (Thermo Fisher Scientific) and imaged on a G:BOX Chemi XX6 gel imager (Syngene). Image J software (National Institutes of Health) was used for densitometry band quantification.

**Supplemental Table 1. Flow cytometry antibody list**

| Antibodies | Source | Identifier |
|---|---|---|
| Brilliant Violet 421 anti-mouse CD117 (c-Kit), Clone 2B8 | BioLegend | Cat# 105828; RRID: AB_11204256 |
| APC/Cyanine7 anti-mouse Ly-6A/E (Sca-1), Clone D7 | BioLegend | Cat# 108126; RRID: AB_10645327 |
| FITC anti-mouse Ly-6A/E (Sca-1), Clone D7 | BioLegend | Cat# 108106; RRID: AB_313343 |
| PE/Cyanine7 anti-mouse CD133, Clone 315-2C11 | BioLegend | Cat# 141210; RRID: AB_2564069 |
| Alexa Fluor 647 anti-mouse CD34, Clone RAM34 | BD Biosciences | Cat# 560230; RRID: AB_1645200 |
| FITC anti-mouse CD34, Clone RAM34 | BD Biosciences | Cat# 553733; RRID: AB_395017 |
| FITC anti-mouse CD45.2, Clone 104 | BD Biosciences | Cat# 553772; RRID: AB_395041 |
| Brilliant Violet 421 anti-human CD117 (c-kit), Clone 104D2 | BioLegend | Cat# 313216; RRID: AB_11148721 |
| Alexa Fluor 647 anti-human CD34, Clone 561 | BioLegend | Cat# 34361; RRID: AB_2632632 |
| PE anti-human CD133, Clone 7 | BioLegend | Cat# 372804; RRID: AB_2632880 |

**Supplemental Table 2. List of TaqMan probes for qRT-PCR.**

| **Probes and primers** | **Identifier** |
|---|---|
| Human 18S | Hs99999901_s1 |
| Mouse Nos2 | Mm00440502_m1 |
| Mouse Il10 | Mm01288386_m1 |
| Mouse Irg1 | Mm01224532_m1 |
| Mouse Nfe2l2 | Mm00477784_m1 |
| Mouse Nqo1 | Mm01253561_m1 |
| Mouse Gata1 | Mm01352636_m1 |
| Mouse EpoR | Mm00833882_m1 |
| Mouse Cpox | Mm00483982_m1 |
| Mouse Gclm | Mm01324400_m1 |
| Mouse Gsr | Mm00439154_m1 |
| Mouse Tnf | Mm00443258_m1 |
| Mouse Hif1a | Mm00458869_m1 |
| Mouse Pdk1 | Mm00554300_m1 |
| Mouse SLC48A1 (Hrg1) | Mm00728070_s1 |
| Mouse β Major Probe | 6FAM- CTCTCTTGGGAACAATTAACCATTGTTCACAG-TAMRA |
| Mouse β Major Forward Primer | 5’ -AACCCCCTTTCCTGCTCTTG- 3’ |
| Mouse β Major Reverse Primer | 5’ -TCATTTTGCCAACAACTGACAGA- 3’ |
| Mouse β H1 Probe | 6FAM- ACTTTCTTGCCATGGGCTCTAATCCGG-TAMRA |
| Mouse β H1 Forward Primer | 5’ -CCTGGCCATCATGGGAAAC- 3’ |
| Mouse β H1 Reverse Primer | 5’-CCCCAAGCCCAAGGATGT-3’ |

**Supplemental Figure 1**

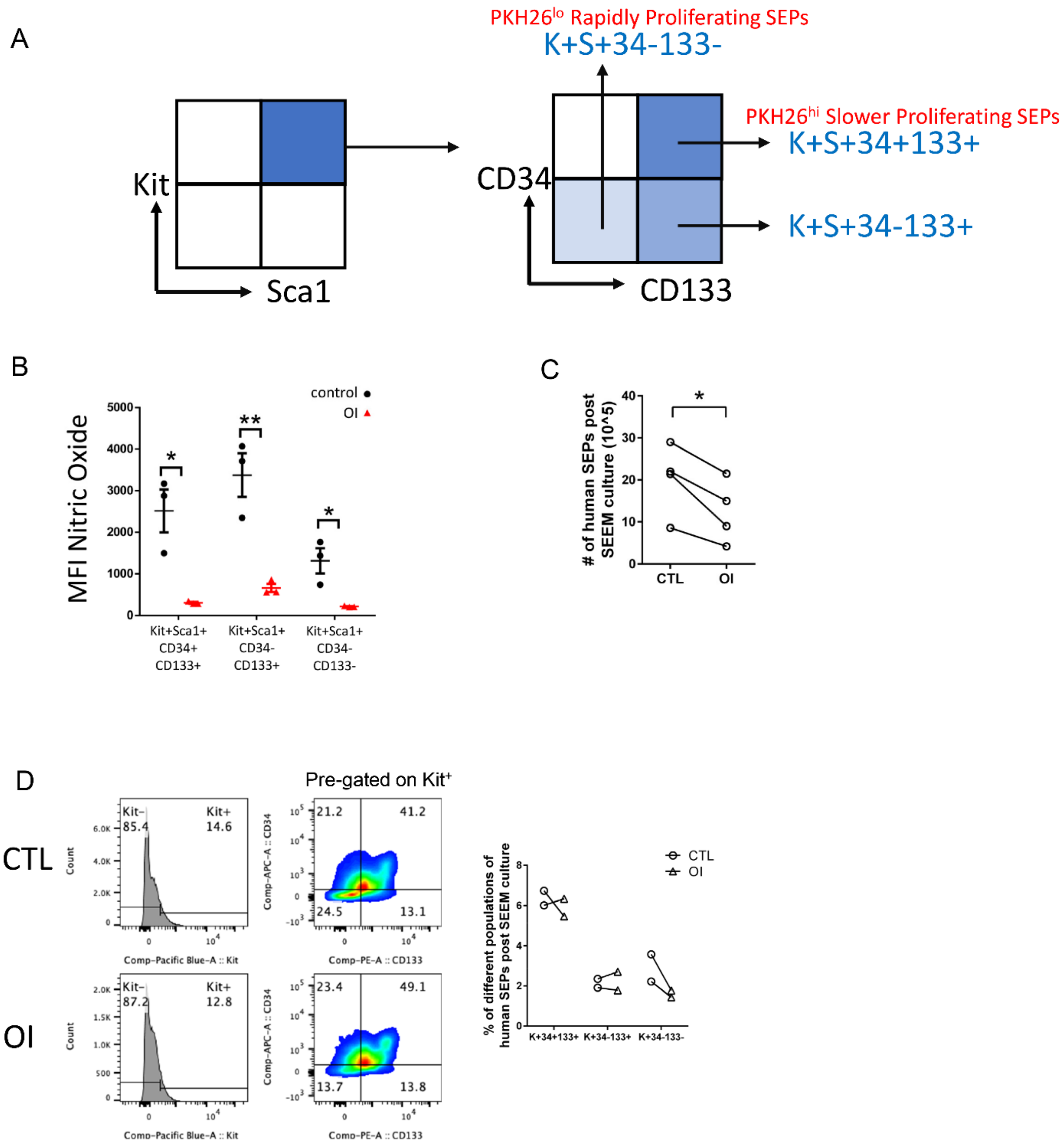

**Supplemental Figure 1. Itaconate blocks SEP expansion by inhibiting iNOS-dependent NO production.**

(A) Gating strategy showing flow cytometry analysis of proliferating SEPs. Cells were stained for viability followed by gating on Kit and Sca1. Pre-gated Kit+Sca1+ cells were then gated on CD34 and CD133 for analysis of different subpopulations. The identification of CD34-CD133-Kit+Sca1+ as rapidly proliferating PKH26$^{lo}$ SEPs and CD34+CD133+Kit+Sca1+ cells as PKH26hi slower proliferating cells comes from ref[(4)].

(B) SEPs were treated ± 125 μM OI at SEEM day 3 for 48 hrs. Quantification of intracellular NO levels different populations of SEPs by mean fluorescence intensity (MFI) of DAF-FM DA staining (E) (n=3 per group, unpaired t test).

(C-D) Human SEEM cultures were treated with vehicle or 125 μM OI for 48 hrs, followed by quantification of total SEP numbers (C), and representative flow cytometry plot (left) and quantification (right) of SEPs (D). Note that Sca1 is not a marker for human SEPs that is why it is not present in this analysis[(1)] (n=4 per group, paired t test (C)).

Data represent mean ± SEM. * $p < 0.05$, ** $p < 0.01$.

**Supplemental Figure 2**

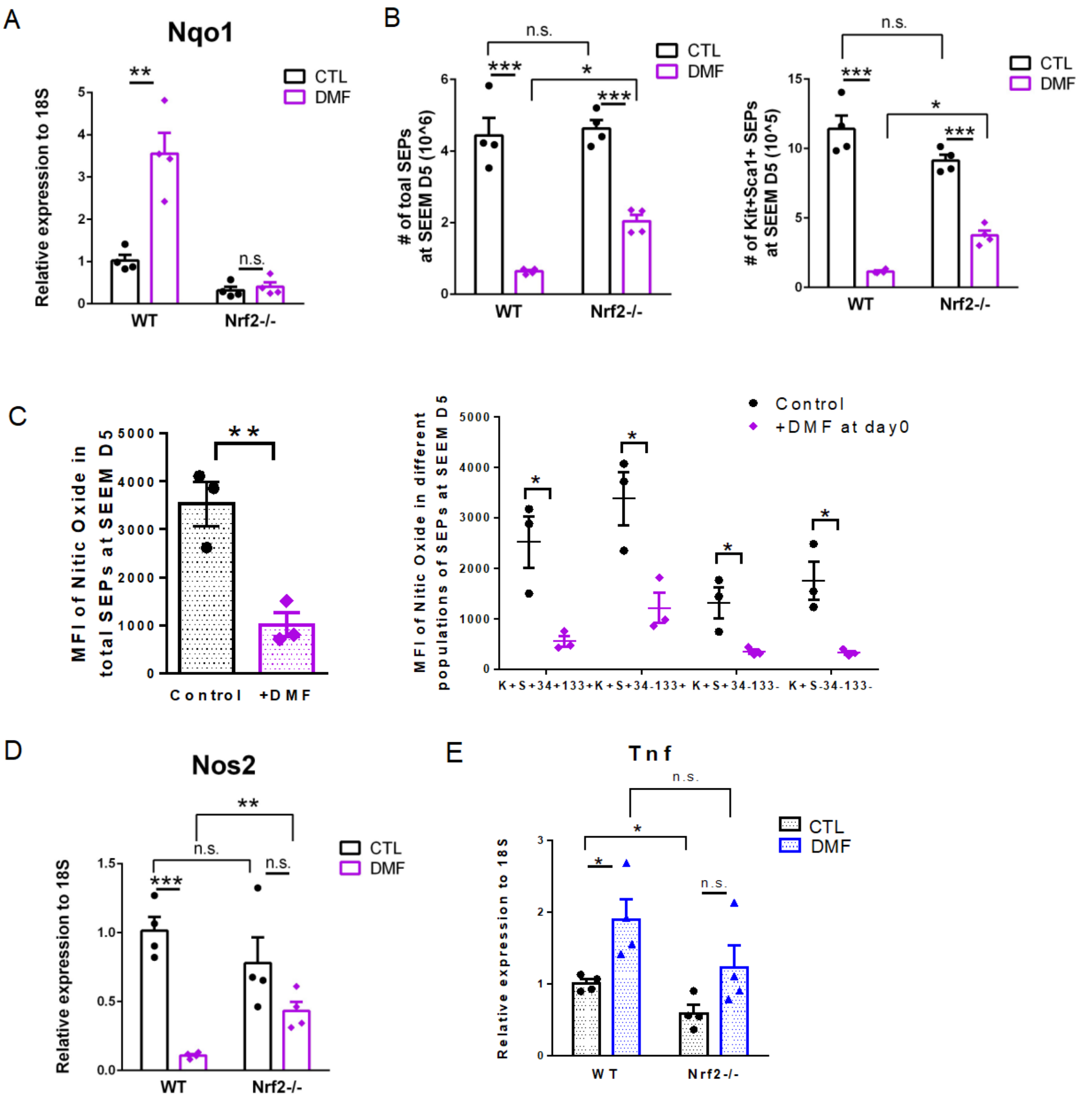

**Supplemental Figure 2. DMF impairs SEP expansion in a Nrf2-dependent manner.**

(A-B) WT and Nrf2-/- SEEM cultures were treated ± 30 μM DMF for 5 days. qRT-PCR analysis of *Nqo1* expression (A), and analysis for numbers of total SEPs (left) and $Kit^{+}Sca1^{+}$ SEPs (right) (B) (n=4; two-way ANOVA/Fisher's LSD).

(C) SEPs were treated ± 30 μM DMF at SEEM day 3 for 48 hrs. Quantification of intracellular NO levels in total SEPs (left) and different SEP populations (right) by MFI of DAF-FM DA staining (E) (n=3 per group, unpaired t test).

(D-E) WT and Nrf2-/- SEEM cultures were treated ± 30 μM DMF for 5 days. qRT-PCR analysis of *Nos2* (D) and *Tnf-α* (E) expression (n=4; two-way ANOVA/Fisher's LSD).

Data represent mean ± SEM. n.s. $p > 0.05$, * $p < 0.05$, ** $p < 0.01$, *** $p < 0.001$.

**Supplemental Figure 3**

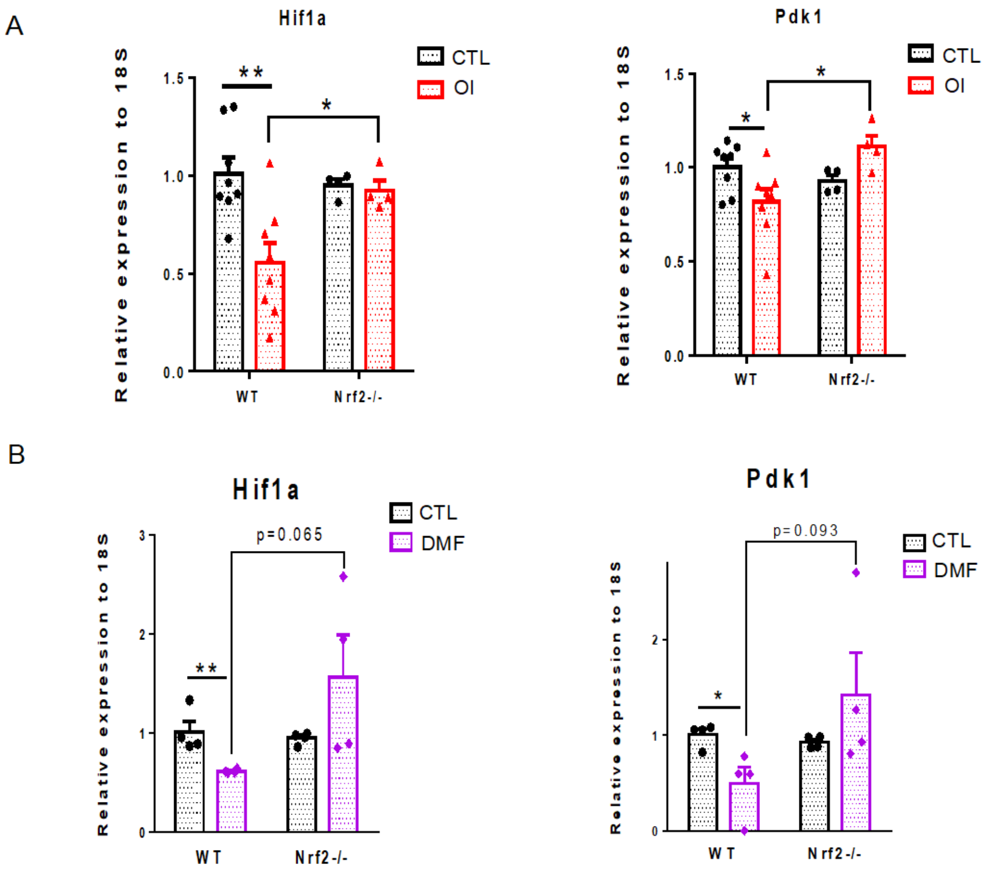

**Supplemental Figure 3. The activation of Nrf2 inhibits Hif-1-dependent glycolysis.**

(A-B) WT and Nrf2-/- SEEM cultures were treated ± 125 μM OI for 5 days. The qRT-PCR analysis of *Hif-1α* (D) and *Pdk1* (E) expression (n=8 in WT and n=4 in Nrf2-/-; two-way ANOVA/Fisher's LSD).

(C-D) WT and Nrf2-/- SEEM cultures were treated ± 30 μM DMF for 5 days. The qRT-PCR analysis of *Hif-1α* (D) and *Pdk1* (E) expression (n=4; two-way ANOVA/Fisher's LSD).

Data represent mean ± SEM. * $p < 0.05$, ** $p < 0.01$.

**Supplemental Figure 4**

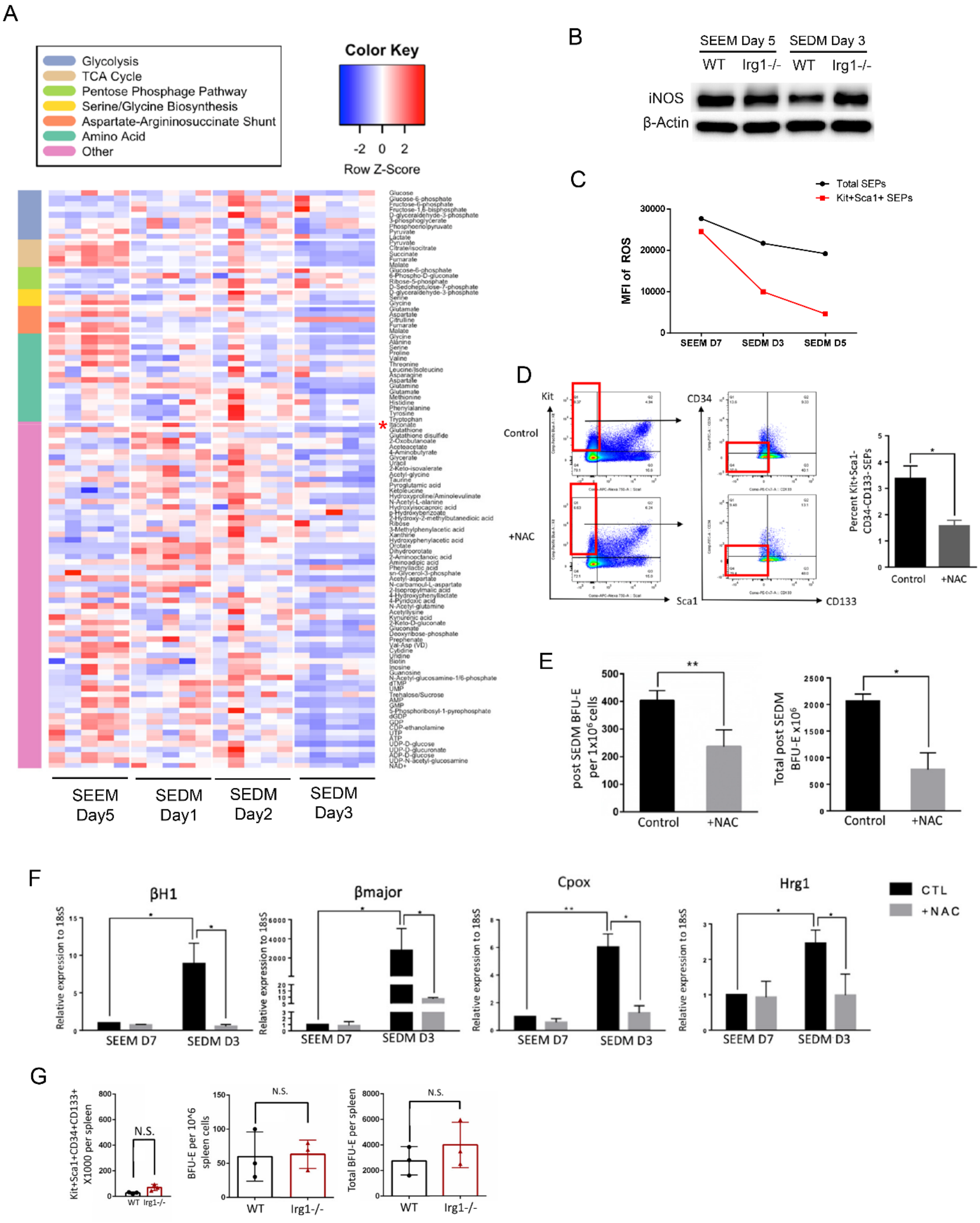

**Supplemental Figure 4. Increased itaconate production couples with the decreased level of ROS during SEP differentiation.**

(A) SEPs were isolated from SEEM and SEDM cultures at indicated days for metabolomics analysis. A heatmap depicting the abundance of metabolites extracted from SEPs in selected pathways. Itaconate is highlighted by asterisk (*). Color represents row-wise scaled z-score of metabolite abundance (n=5 per time points).

(B) SEPs were harvested from WT, or Irg1-/- cultures at SEEM Day 5 and SEDM Day 3. WB analysis of iNOS protein expression.

(C) SEPs isolated from SEEM and SEDM cultures at noted time points were analyzed for intracellular ROS levels in total SEPs and $Kit^{+}Sca1^{+}$ SEPs. ROS levels were quantified by MFI of CellROX staining.

(D-F) Cells were treated with ± 5mM NAC at day 4 of SEEM culture for 24 hours, and cells were switched to SEDM culture for 3 days. Representative flow cytometry plot showing analysis of SEPs at SEDM day3 using Kit, Sca1, CD34 and CD133 as markers (left), and percentage of Kit+Sca1-CD34-CD133- SEPs (right) (D). Frequency (left) and total numbers (right) of stress BFU-E present in the differentiation culture (E). qRT-PCR analysis of erythroid-associated gene expression (F).

(G) Analysis of CD34+CD133+Kit+Sca1+ SEPs and stress BFU-E in wildtype and Irg1-/- spleens prior to treatment with HKBA. Total number of CD34+CD133+Kit+Sca1+ SEPs in the spleen as analyzed by flow cytometry and total cellularity counts (left). Frequency of stress BFU-E (middle) and total number of stress BFU-E(right) in the spleen prior to HKBA treatment (n=3 per group).

Data represent mean ± SEM. N.S. $p > 0.05$, * $p < 0.05$, ** $p < 0.01$.

### Supplemental Figure 5

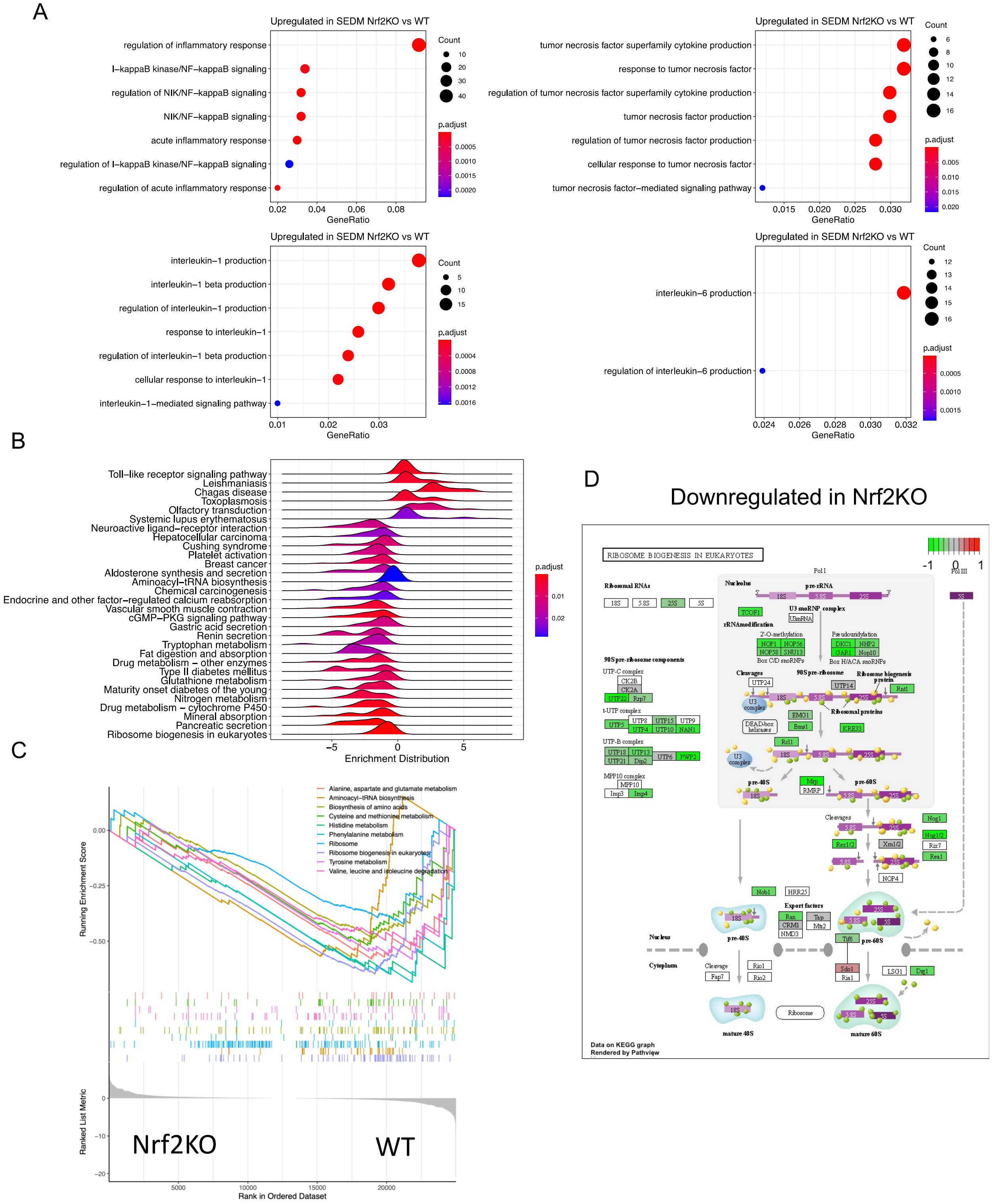

**Supplemental Figure 5. RNA-seq analysis comparing WT and Nrf2-/- SEPs isolated from SEDM cultures.**

(A) Overrepresentation analysis of upregulated DEGs in SEDM Nrf2-/- (FDR < 0.05, FC of Nrf2-/-/WT > 1.5) showing significant enriched GO terms related to inflammatory response, NF-κB pathway and inflammatory cytokine-dependent signaling. Circle size represents the numbers of genes in each GO term and color represents adjusted p value (n=3).

(B) GSEA analysis of top enriched KEGG pathways between WT and Nrf2-/- differentiating SEPs (n=3).

(C) GSEA analysis of AA- and ribosome-associated GO terms.

(D) KEGG pathviewer map of ribosome biogenesis in eukaryotes.

### Supplemental Figure 6

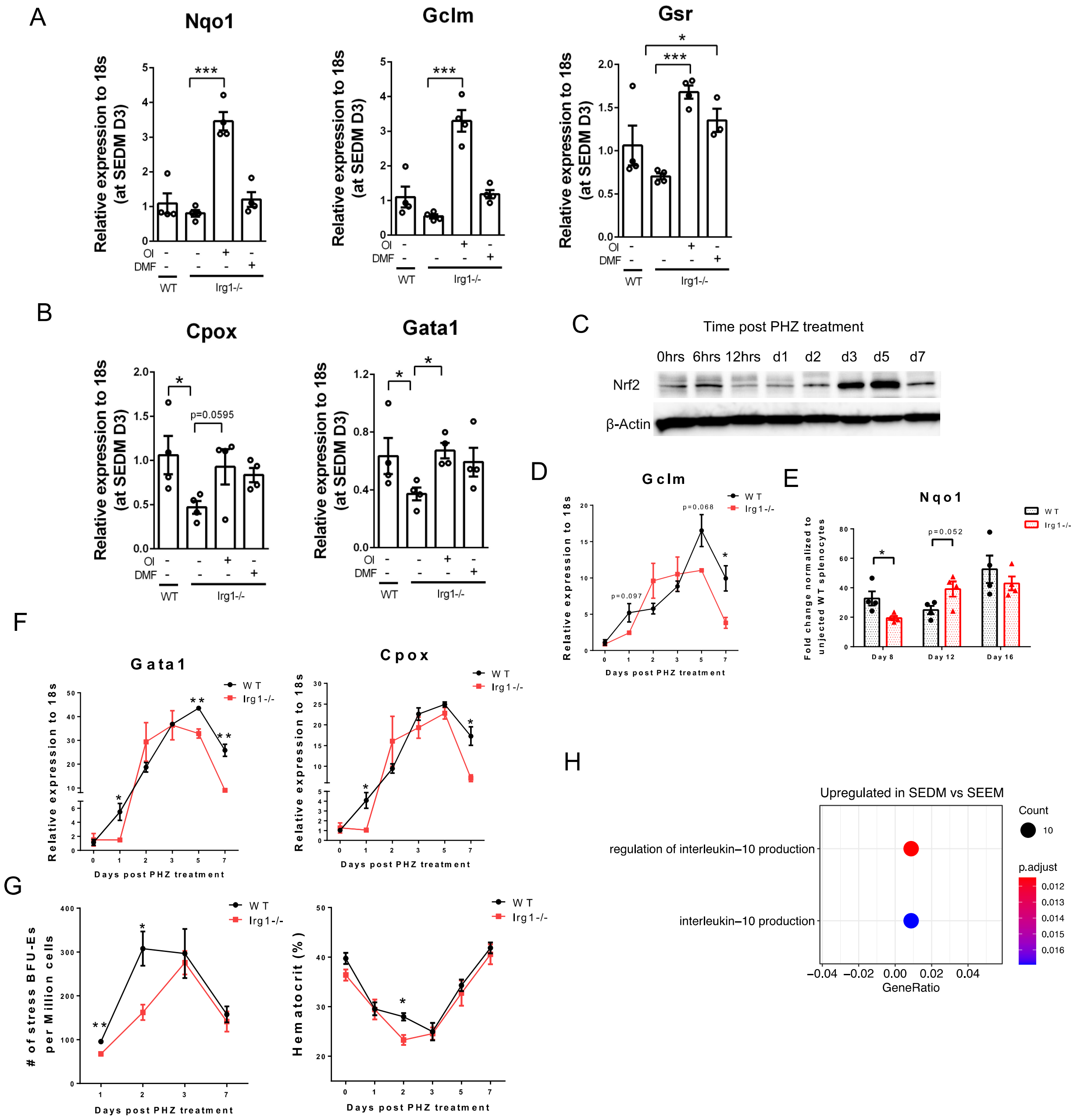

**Supplemental Figure 6. Itaconate promotes Nrf2-dependent SEP differentiation.**

(A-B) SEPs were harvested from WT, or Irg1-/- SEDM cultures treated with vehicle, 125 μM OI or 30 μM DMF for 3 days. Analysis of mRNA expression of Nrf2 target genes (A) and erythroid-specific genes (B) by qRT-PCR (n=4 per group, one-way ANOVA/Tukey's).

(C) WT mice were injected intraperitoneally with a single dose (100 mg/kg body weight) of freshly prepared phenylhydrazine. WB analysis of Nrf2 protein levels at indicated time point post PHZ injection.

(D) Age- and sex-matched WT and Irg1-/- mice were injected intraperitoneally with a single dose (100 mg/kg body weight) of freshly prepared phenylhydrazine. qRT-PCR detection of mRNA levels of *Gclm* (n=3 per time point, unpaired t test).

(E) Age- and sex-matched WT and Irg1-/- mice were administered with HKBA ($5 \times 10^8$ particles/mouse) via intraperitoneal injection. qRT-PCR detection of mRNA levels of *Nqo1* (n=4 per time point, unpaired t test).

(F-G) Experiment was described in (D). qRT-PCR detection of mRNA levels of erythroid-specific genes (F). Frequency of BFU-Es (left) and hematocrit level (right) at indicated time point post PHZ injection (G) (n=3 per time point, unpaired t test).

(H) RNA-seq analysis comparing SEPs isolated from SEEM and SEDM cultures. Overrepresentation analysis of upregulated DEGs in SEDM (FDR < 0.05, Fold change (FC) of SEDM/SEEM > 1.5), showing GO terms related to IL-10 production. Circle size represents the numbers of genes in each pathway and color represents BH-adjusted p value (n=3 per group).

Data represent mean ± SEM. * $p < 0.05$, ** $p < 0.01$, *** $p < 0.001$.

### Supplemental Figure 7

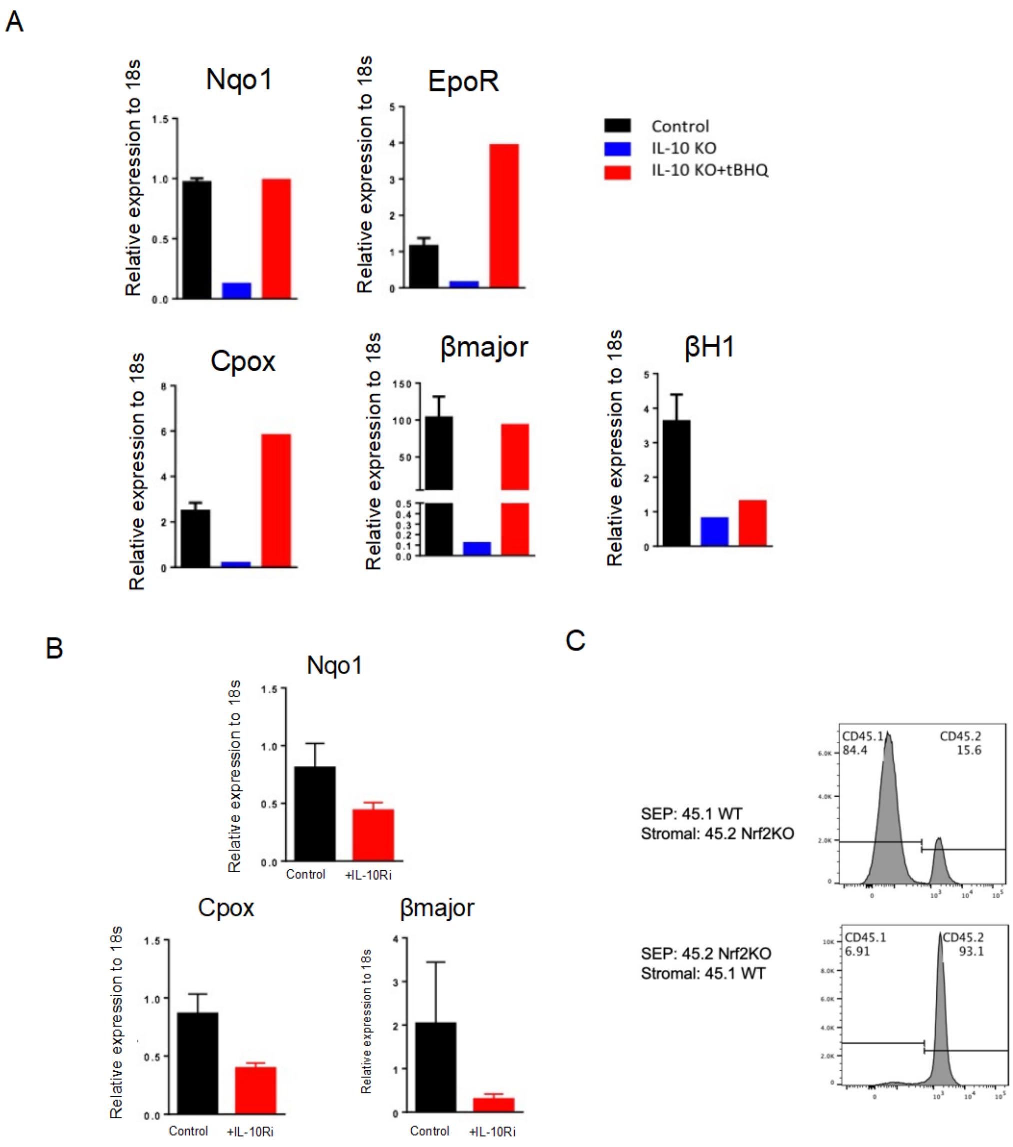

**Supplemental Figure 7. IL-10 promotes Nrf2-dependent SEP differentiation.**

(A) WT and IL10-/- BM cells were placed into SEEM for 7 days, followed by differentiation in SEDM for 3 days. IL10-/- SEDM cultures were treated ± 20μM tBHQ. Analysis of mRNA expression of Nrf2 target genes and erythroid-specific genes by qRT-PCR detection.

(B) WT BM cells were cultured in SEEM for 7 days, followed by differentiation in SEDM treated ± IL-10 receptor alpha blocking antibody. Analysis of mRNA expression of Nrf2 target genes and erythroid-specific genes by qRT-PCR detection.

(C) WT (CD45.1) and Nrf2-/- (CD45.2) BM cells were cultured in SEEM for 5 days followed by 3 days in SEDM. When switched to SEDM, nonadherent SEPs were collected and plated on the stromal layer from indicated genotype. SEPs isolated from SEDM cultures were subject to flow cytometry analysis by staining CD45.2.

Data represent mean ± SEM.

### Supplemental data references


1. Xiang J, Wu DC, Chen Y, Paulson RF. In vitro culture of stress erythroid progenitors identifies distinct progenitor populations and analogous human progenitors. Blood. 2015;125(11):1803-12. Epub 2015/01/23. doi: 10.1182/blood-2014-07-591453. PubMed PMID: 25608563; PMCID: 4357585.

2. Bennett LF, Liao C, Paulson RF. Stress Erythropoiesis Model Systems. Methods Mol Biol. 2018;1698:91-102. Epub 2017/10/28. doi: 10.1007/978-1-4939-7428-3_5. PubMed PMID: 29076085; PMCID: PMC6510234.

3. Gardenghi S, Renaud TM, Meloni A, Casu C, Crielaard BJ, Bystrom LM, Greenberg-Kushnir N, Sasu BJ, Cooke KS, Rivella S. Distinct roles for hepcidin and interleukin-6 in the recovery from anemia in mice injected with heat-killed Brucella abortus. Blood. 2014;123(8):1137-45. Epub 2013/12/21. doi: 10.1182/blood-2013-08-521625. PubMed PMID: 24357729; PMCID: PMC3931188.

4. Hao S, Xiang J, Wu DC, Fraser JW, Ruan B, Cai J, Patterson AD, Lai ZC, Paulson RF. Gdf15 regulates murine stress erythroid progenitor proliferation and the development of the stress erythropoiesis niche. Blood Adv. 2019;3(14):2205-17. Epub 2019/07/22. doi: 10.1182/bloodadvances.2019000375. PubMed PMID: 31324641; PMCID: PMC6650738 interests.
